# Antagonistic histone H2A variants and autonomous heterochromatin formation shape epigenomic patterns in Arabidopsis

**DOI:** 10.1101/2025.10.19.683276

**Authors:** Shoko Oda, Sayaka Tominaga, Shumpei Takeuchi, Akihisa Osakabe, Akira Kawabe, Frédéric Berger, Tetsuji Kakutani, Taiko Kim To

**Affiliations:** Department of Biological Sciences, The University of Tokyo, Tokyo 113-0033, Japan; School of Life Science and Technology, Institute of Science Tokyo, Kanagawa 226-8501, Japan; Graduate School of Science, Chiba University, Chiba 263-8522, Japan; Institute for Advanced Academic Research, Chiba University, Chiba 263-8522, Japan; Faculty of Life Sciences, Kyoto Sangyo University, Kyoto 603-8555, Japan; Gregor Mendel Institute (GMI), Austrian Academy of Sciences, Vienna Biocenter (VBC), Vienna, Austria

## Abstract

Heterochromatin formation is pivotal in many eukaryotes with repetitive sequences, such as transposable elements (TEs). However, in plants, where the known *de novo* DNA methylation mechanism (RdDM) targets euchromatin, how heterochromatin is formed in a region-specific manner remains unclear. We previously reported an RdDM-independent *de novo* establishment of H3K9me and non-CpG methylation, both of which localize in heterochromatin. Here we show that the mutually exclusive histone H2A variants, H2A.W and H2A.Z, function as guides to initiate heterochromatin formation; H2A.W and H2A.Z promotes and inhibits heterochromatin establishment, respectively, especially in chromosomal arm regions with dispersed TEs. In contrast, pericentromeric TEs demonstrate autonomous heterochromatin formation, less dependently on these H2A variants. Furthermore, H2A.Z protects protein-coding genes from ectopic heterochromatin formation, possibly by preventing its spreading. We propose that the genome indexing mechanism driven by H2A variants, as well as the autonomous formation of pericentromeric heterochromatin, shapes proper epigenomic patterns in Arabidopsis.

## Introduction

Most eukaryotic genomes contain a substantial amount of repetitive elements and transposable elements (TEs). TEs are abundant in pericentromeric heterochromatin, and are silenced by epigenetic modifications, including the methylation of cytosine (mC) and histone H3 lysine 9 (H3K9me)^1,2^. In plants, mC can be classified into those in CpG sites (mCG) and non-CpG sites (mCH, H = A, T, C)^3,4^. mCH colocalizes with H3K9me in TEs, whereas mCG is found in both TEs and bodies of active genes^5–7^. It is crucial for plants to establish and maintain proper distribution of epigenetic information in the genome (epigenome patterns), and to silence TEs specifically^8,9^. However, in contrast to the intensive studies of vertebrates which involve sequence-driven heterochromatin formation^1,10^, the mechanism by which hosts distinguish TEs from genes and initiate heterochromatin formation specifically at TEs has long remained obscure in plants.

Pathways controlling heterochromatin marks have been extensively investigated in Arabidopsis. H3K9me is catalyzed by Suv39H-homolog histone methyltransferases, SUVH4, SUVH5, and SUVH6^11–13^. mCH is further classified into mCHG and mCHH, and is mostly catalyzed by plant-specific CpH methyltransferases, CHROMOMETHYLASE 3 (CMT3) and CMT2, respectively^14–17^. H3K9me and mCH is maintained by a positive feedback between them, while H3K9me/mCH and mCG are maintained almost independently of each other^11,12,18,19^. mCG is maintained by METHYLTRANSFERASE 1 (MET1), a homolog of mammalian DNMT1^20,21^. Unlike in mammals, the *de novo* DNA methyltransferase DRM2 is recruited in plants via RNA interference (RNAi) machinery in a process called RNA-directed DNA methylation (RdDM)^9,22,23^. However, unlike in fission yeast, where RNAi machinery recruits the Suv39H homolog Clr4 to pericentric heterochromatin^24^, DRM2 is guided to short non-coding TEs in euchromatic chromosome arms. How pericentromeric heterochromatin is formed remains unclear.

We previously reported identification of a novel pathway for establishing *de novo* mCH/H3K9me specifically to TEs^25,26^. This pathway is independent of RdDM and suggested to involve CMTs and SUVHs. It predominantly functions in long TEs, most of which are in pericentromeric heterochromatin. However, how they specifically target TEs is still enigmatic.

Genomic distribution of TEs and genes across chromosomes is associated with that of histone H2A variants. H2A.Z, an evolutionarily well-conserved variant, is localized to protein-coding genes in euchromatic chromosome arms^27–29^, whereas a flowering plant-specific H2A variant, H2A.W, is specifically localized in TEs, and enriched in constitutive heterochromatin in pericentromeric regions^29–31^. Recently, DDM1, which has a significant impact on the maintenance of DNA methylation in TEs, binds and deposits H2A.W over TEs and potentially removes H2A.Z at TEs^29,32–34^. However, DNA methylation is only moderately decreased in *h2a.w* heterochromatin^31^, and rarely increases at TEs in *h2a.z*^28^, obscuring their roles. In addition, H2A.W and H3K9 methylation jointly suppress expression of TEs, indicating a major role of this variant in silencing^35^.

Here, we directly investigate the role of H2A.W and H2A.Z in heterochromatin formation. Results from mCH reconstitution in the absence of H2A variants reveal that H2A.W can promote mCH establishment in TEs, while H2A.Z can suppress it in TEs and protein-coding genes. These effects are particularly evident in TE-poor chromosomal arms. In these regions, when mCH is lost, H2A.W is replaced by H2A.Z at hundreds of TEs, attenuating subsequent mCH establishment. In contrast, this replacement is not observed in pericentromeric TEs where mCH efficiently recovers. However, when both mCG and mCH are lost, pericentromeric H2A.W is also replaced by H2A.Z. The resulting simultaneous loss of H2A.W, mCG and mCH can all be reversed by complementation of the mCH machinery, but this “autonomous” recovery of heterochromatin marks is only seen in TE-rich pericentromeric regions. These contrasting dynamics of heterochromatin formation between TE-dense pericentromeric and TE-poor arm regions both account for the proper differentiation of epigenome patterns in Arabidopsis.

## Results

### H2A.W is required for mCH establishment in hundreds of TEs

Previously, we developed the mCH/H3K9me reconstitution system to investigate the dynamics and mechanism of mCH/H3K9me establishment genome-wide^25^. In the system, two mCH/H3K9me-deficient mutants (*cmt2 cmt3* (*cc*) and *suvh4 suvh5 suvh6* (*sss*)) were genetically crossed to produce F1 plants. F1 plants restore all *CMTs* and *SUVHs* genes as heterozygous (Fig. 1a, left), allowing us to assess the dynamics of mCH/H3K9me recovery. As shown previously, F1 plants restore both mCH/H3K9me at most TE genes (coding regions within TEs; hereafter referred to as TEs). More than 96% of TEs (3419 out of 3538 TEs with mCHG in wild-type (WT) >0.1; 3389 out of 3471 TEs with mCHH in WT >0.03) show efficient mCH recovery, while they fail to recover mCH/H3K9me at a subset of TEs, which we previously designated as Gene-Like TEs (GLTs; n=73, F1/WT <0.1 for mCHG)^25^. The failure in GLTs is suggested to involve histone H2A variants, because the replacement from heterochromatic H2A.W to euchromatic H2A.Z is specifically observed at GLTs. These led us to investigate the impacts of histone H2A variants on mCH establishment.

**Fig. 1.**
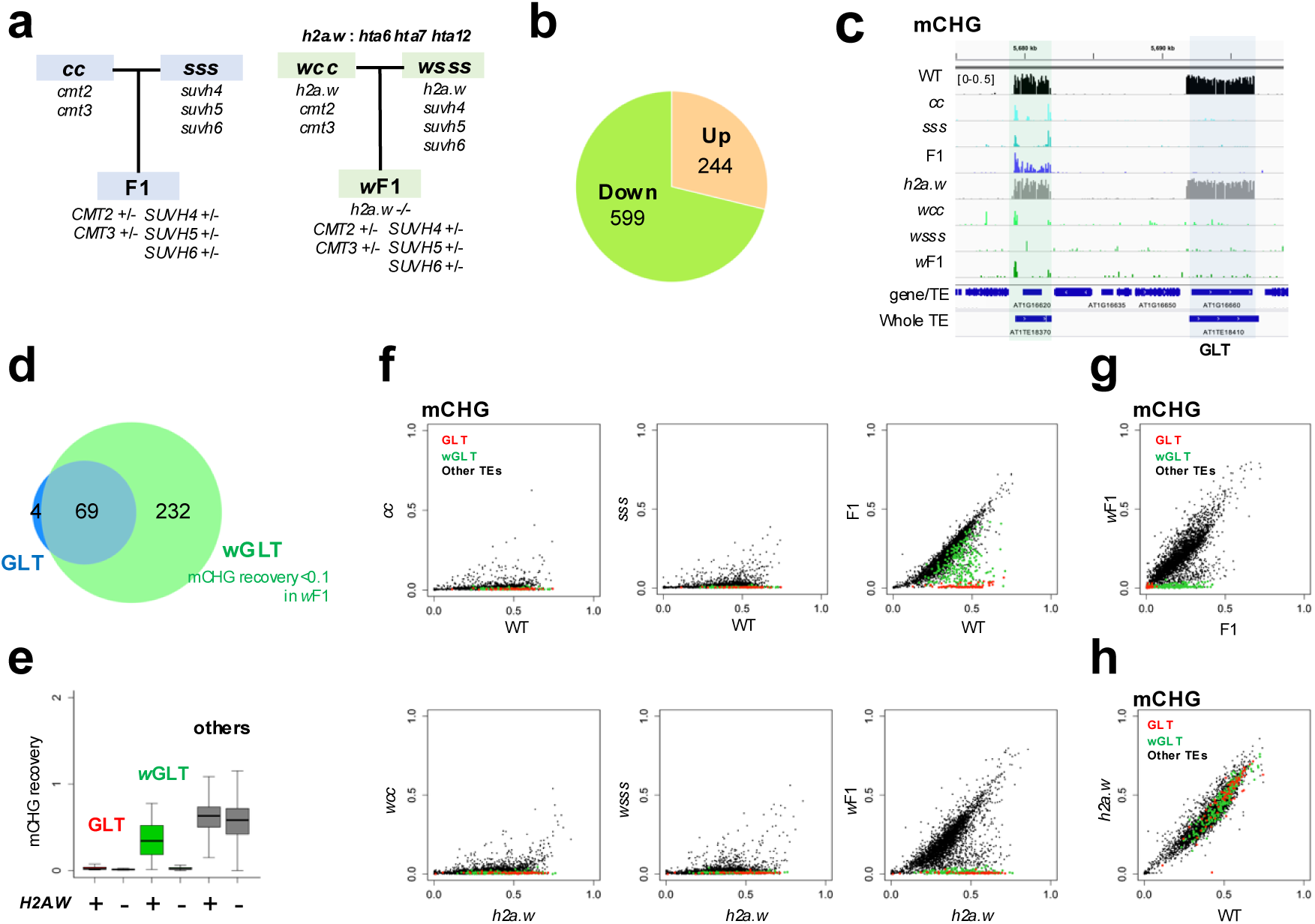
*h2a.w* mutation reduces the efficiency of mCH recovery in hundreds of TEs. **a** Scheme of mCH/H3K9me reconstitution in *H2A.W* (left) and in *h2a.w* (right) background. The mutant of CpH methyltransferases (*cmt2 cmt3*: *cc*) and the mutant of H3K9 methyltransferases ( *suvh4 suvh5 suvh6*: *sss*) were crossed to create F1 plants heterozygous for all the mutated genes (WGBS data from GSE148753; To et al., 2020). The *h2a.w cc* mutant (*hta6 hta7 hta12 cmt2 cmt3*: *wcc*) and the *h2a.w sss* mutant (*hta6 hta7 hta12 suvh4 suvh5 suvh6*: *wsss*) were crossed to create F1 plants in *h2a.w* mutation background (*w*F1). **b** Pie chart showing the number of TEs with statistically significant changes in m CHG in *w*F1 compared to the original F1. **c** Genome browser view of mCHG in the indicated plants. TE with recovery in F1 but not in *w*F1 is highlighted in green. TE without recovery in both F1 and *w*F1 is highlighted in blue. The region Chr1/5,675,000–5,698,000 is shown. **d** Venn diagram showing the overlap between GLTs (blue) and TEs without recovery in *w*F1 (green). TEs with recovery in F1 but not in *w*F1 are designated as wGLTs (n=232). **e** Comparison of mCHG recovery between F1 and *w*F1. The recovery was calculated as F1/WT (H2A.W +) and *w*F1*/h2a.w* (H2A.W-), respectively. To avoid division by values near zero, TEs with mCHG (>0.1) in both WT and *h2a.w* were analyzed (n=3,511). Outliers are not shown. The centerline and box edges represent quartiles, and whiskers range 1.5 times of the interquartile from the box edges. **f** mCHG level for each TE in *cc*, *sss*, and F1 compared to WT, as well as *wcc*, *wsss*, and *w*F1 compared to *h2a.w*. GLTs are in red, wGLTs are in green, and the other TEs are in black. **g** Comparison between F1 and *w*F1 of mCHG levels for each TE. **h** Comparison between WT and *h2a.w* of mCHG levels for each TE.

We first introduced *h2a.w-2* (*hta6 hta7 hta12*) mutation^31^ into the mCH/H3K9me reconstitution system, and examined the impacts of H2A.W on mCH recovery (Fig. 1a, right). The parental mutants, *h2a.w cc* (*wcc*) and *h2a.w sss* (*wsss*), as well as their F1 progeny (*w*F1), were subjected to BS-seq analysis. The results were then compared with those of the original F1 (GSE148753^25^). Indeed, *h2a.w* mutation significantly altered mCH recovery (Fig. 1b-g, S1a-f); statistically significant changes in mCHG and mCHH levels were observed at 843 TEs and 1067 TEs in *w*F1 compared to the original F1 (Fig. 1b,c, S1b,c), respectively. Most of them exhibited hypomethylation in *w*F1 (mCHG 71.1%; mCHH 84.2%) (Fig. 1b, S1b). Importantly, as previously reported^31^, *h2a.w* mutation alone negligibly affected mCH maintenance (Fig. 1h, S1g), demonstrating the contrasting effects of H2A.W on the maintenance and establishment.

The *h2a.w* mutation increased the number of TEs without recovery by approximately four times in *w*F1 (301 TEs; *w*F1/*h2a.w* <0.1 for mCHG) (Fig. 1d), compared with those in the original F1 (73 GLTs; F1/WT <0.1 for mCHG). These include most GLTs (69 out of 73) and additional 232 TEs; hereafter the latter called wGLTs. wGLTs require H2A.W for the mCH establishment (Fig. 1c-g), while mCH recovery in wGLTs was incomplete even in F1 harboring the wild-type *H2A.W* genes (Fig. 1e,f, S1d,e, S2). The *h2a.w* mutation also suppressed the mCH recovery in a number of TE fragments and TE-like genes with methylation (mCHG in WT>0.1) (Fig. S3). These results indicate that H2A.W can enhance mCH establishment in hundreds of TEs, TE fragments, and TE-like genes.

### *h2a.z* mutation suppresses the failure of mCH establishment in GLTs and wGLTs

In our previous study^25^, the failure of mCH/H3K9me recovery at GLTs in F1 was associated not only with H2A.W loss but also with H2A.Z gain, suggesting that H2A.Z might also have an impact on mCH/H3K9me establishment. To test this possibility, we tried to create a F1 in *h2a.z* (*hta8 hta9 hta11*) background by crossing *h2a.z cc* (*zcc*) and *h2a.z sss* (*zsss*). However, *zsss* exhibited too severe developmental defects including infertility to use for crossing. By contrast, *zcc* mutant was slightly fertile while having global mCH loss (Fig. S4a), which allowed us to investigate mCH recovery dynamics in the absence of H2A.Z, by complementing *CMT* genes via transformation. As controls, we firstly created T1 plants by transforming *CMT3* and *CMT2* transgene, separately, into *cc* (hereafter called *cc*+*CMT3* and *cc*+*CMT2*, respectively, Fig. 2a, S5a). mCH recovery was then examined by BS-seq. In the two individual *cc*+*CMT3*s, mCHG was consistently recovered substantially in most TEs (Fig. 2b) as wild-type levels, while mCHH did not largely recover (Fig. S4b other TEs), which agrees with predominant mCHG activity of CMT3^14–17^. Importantly, in agreement with the observation in F1, mCHG recovery was hardly observed in GLTs and was attenuated in wGLTs (Fig. 2b,c). In the two individual *cc*+*CMT2*s, mCHH recovered substantially for most TEs (Fig. S5b-e). This mCHH recovery was compromised in GLTs and attenuated in wGLTs (Fig. S5b–d). These results indicate that the respective complementation of *CMT3* and *CMT2* genes into *cc* can mimic the mCH/H3K9me reconstitution system using F1 hybrids.

**Fig. 2.**
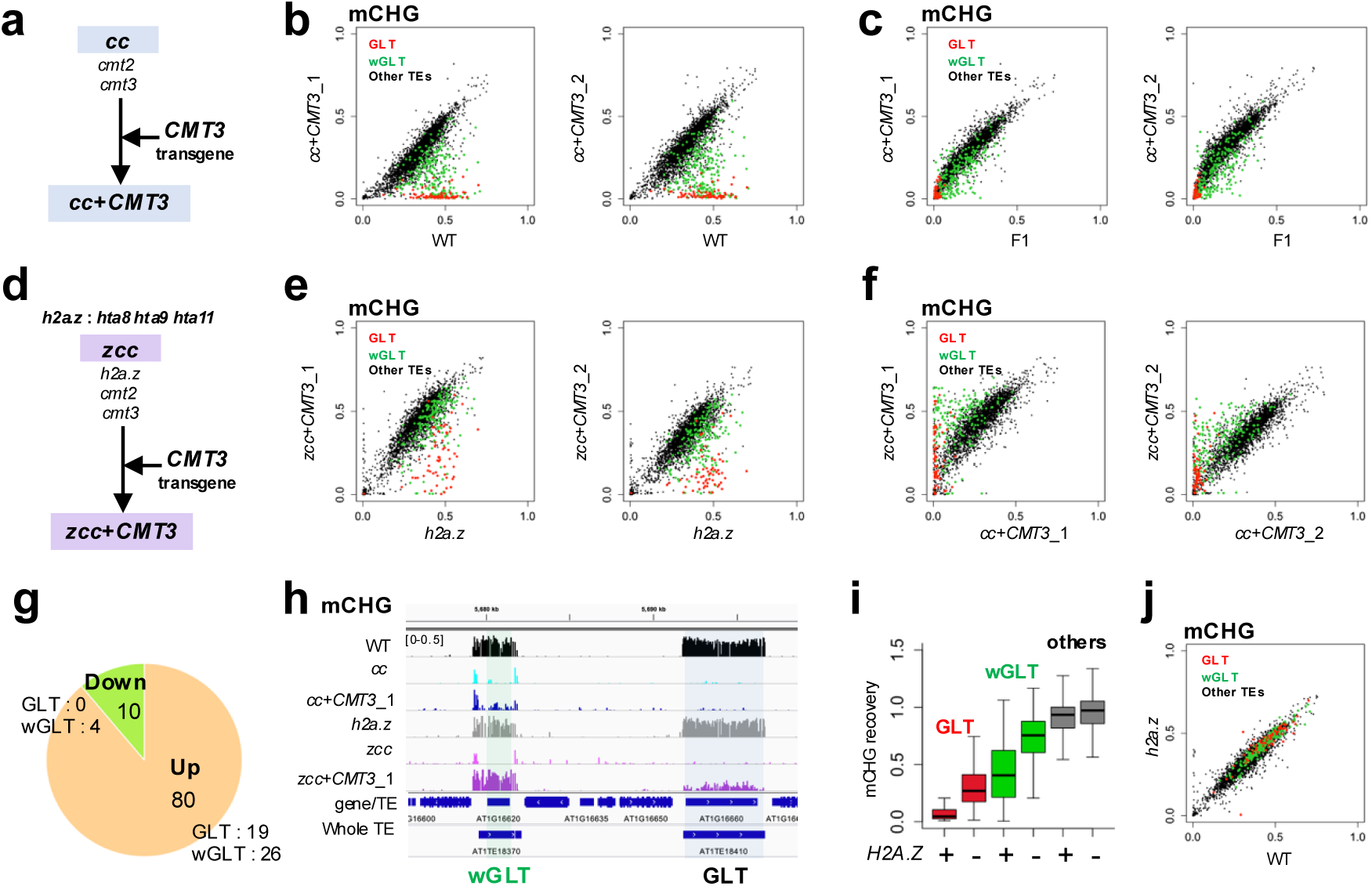
*h2a.z* mutation enables mCH recovery in GLTs and wGLTs. **a,d** Scheme of mCH/H3K9me reconstitution in *H2A.Z* (**a**) and in *h2a.z* (**d**) background. *CMT3* gene was complemented by transforming the *CMT3* transgene with its native promoter into *cmt2 cmt3* (*cc*; **a**) and into *hta8 hta9 hta11 cmt2 cmt3* (*zcc*; **d**), respectively. **b,c** Comparison of mCHG levels for each TE in *cc*+*CMT3* individuals versus WT (**b**) and F1 (**c**). Data for WT and F1 are from GSE148753. GLTs are in red, *w*GLTs are in green, and the other TEs are in black. **e,f** Comparison of mCHG levels for each TE in *zcc+CMT3* individuals versus *h2a.z* (**e**) and *cc*+*CMT3* (**f**). **g** Pie chart showing the number of TEs with statistically significant changes in mCHG between three *zcc*+*CMT3*s (1-3) compared to that in two *cc*+*CMT3s*. **h** Genome browser view of mCHG as in the format of Fig. 1**c**. **i** Comparison of mCHG recovery in *cc*+*CMT3s* and *zcc*+*CMT3*s. Recovery was calculated as the mean mCHG levels of two *cc*+*CMT3*s to WT (*H2A.Z* +) and the mean mCHG levels of three *zcc*+*CMT3*s to *h2a.z* (*H2A.Z*-), respectively. To avoid division by values near zero, TEs with mCHG (>0.1) in both WT and *h2a.z* were analyzed (n = 2,904). Outliers are not shown. The centerline and box edges represent quartiles, and whiskers range 1.5 times of the interquartile from the box edges. **j** Comparison of mCHG levels for each TE between WT and *h2a.z*.

The same strategy was taken to examine the impact of H2A.Z on mCH recovery. To complement *CMT* genes in *h2a.z* background, we transformed *CMT3* and *CMT2*-transgene separately into *zcc* mutant (hereafter *zcc*+*CMT3* and *zcc*+*CMT2* respectively)(Fig. 2d, S5f). mC was profiled for each of six and three individual T1 plants (*z*T1s), respectively, as well as their non-transgenic controls. Indeed, *h2a.z* mutation significantly affected mCH establishment dynamics (Fig. 2e–i, S4c,d, S5h-j). Many TEs showed significantly altered mCHG recovery in *zcc*+*CMT3*s, as well as mCHH in *zcc*+*CMT2*s, in comparison to their respective controls with functional H2A.Z (Fig. 2g, S5g). Most of them exhibited hypermethylation, indicating that *h2a.z* mutation enhances mCH recovery in TEs. Strikingly, *h2a.z* mutation enabled the mCH recovery in GLTs and wGLTs, with substantial mCH recovery being observed in *z*T1s (Fig. 2e–i, S4c,d, S5h,i). In order to see if the enhanced mCH recovery was due to enhanced expression of *CMT3* and *CMT2* transgenes, we examined their transcript levels. Generally, *CMT3* and *CMT2* transcript levels in *z*T1s were mostly comparable to or slightly lower than that in WT and F1 (Fig S4f, S5k,l), implying that enhanced mCH recovery in *h2a.z* background was not due to overexpression of *CMT* transgenes (see legends of Figs S4f, S5k-m for detail). These results suggest that ectopically increased H2A.Z can cause the failure of mCH recovery in GLTs as well as attenuation in wGLTs. *h2a.z* mutation also significantly enabled mCHG recovery in *zcc*+*CMT3*s and mCHH recovery in *zcc*+*CMT2*s in TE fragments and TE-like genes (Fig. S4g,h, S5n,o). In contrast, *h2a.z* mutation alone did not increase mCH maintenance in most TEs, TE fragments and TE-like genes (Fig. 2j, S4e), as previously reported^28^. Taken together, these results suggest that, despite its negligible effect on mCH maintenance, H2A.Z has a negative impact on mCH establishment in a significant fraction of TEs, TE fragments and TE-like genes.

### H2A.W and H2A.Z mutually inhibit their accumulation in *CMTs/SUVHs* mutants

The results so far demonstrate that *h2a.w* and *h2a.z* mutations have opposite effects on mCH establishment. The opposite behavior was also observed in terms of their localization in GLTs; H2A.W decrease and H2A.Z increase in *cc*, *sss* and F1^25^. Therefore, we next asked whether *h2a.w* mutation impacts H2A.Z localization in mCH mutants, and vice versa, by performing chromatin immunoprecipitation-sequencing (ChIP-seq).

First, we analyzed their localization in WT, *cc*, *sss*, and F1. A decrease in H2A.W and an increase in H2A.Z compared to WT were observed, not only at GLTs but also at wGLTs (Fig. 3a-f, S6). Their change levels were negatively correlated with mCH recovery in F1 (i.e. the highest in GLTs, then wGLTs, but barely in the other TEs, while mCH recovery shows the opposite trend). Then, we tested whether *h2a.w* mutation causes an increase in ectopic H2A.Z localization particularly in wGLTs, where the mCH recovery was lost in *h2a.w* background whereas it is enhanced in *h2a.z* background. Although *h2a.w* mutation alone rarely changed H2A.Z distribution^31,36^ (Fig. 3a-c, S6a), it did increase H2A.Z occupancy in wGLTs in *wcc*, *wsss*, and *w*F1 compared to those in *cc*, *sss*, and F1, particularly around TSS and TTS (Fig. 3a-c, S6a,b). The change in H2A.Z was negatively correlated with mCG as observed previously^27^, and TEs exhibiting both ectopic H2A.Z and mCG loss showed inefficient mCH recovery in F1 and *w*F1 (Fig. 3g,h). As for GLTs, strong ectopic H2A.Z was already observed in *cc*, *sss* and F1, and *h2a.w* mutation did not cause further increase in H2A.Z occupancy, which could be explained by substantial depletion of H2A.W at GLTs in WT background (Fig. 3a–f, S6). The other TEs showed minimal sensitivity to either *h2a.w* and *cc*/*sss* mutations, with H2A.Z levels remaining at low levels in all the plants examined. These results suggest that H2A.W can antagonize H2A.Z localization to inhibit ectopic H2A.Z at least at wGLTs and possibly at GLTs, only when mCH/H3K9me is absent.

**Fig. 3.**
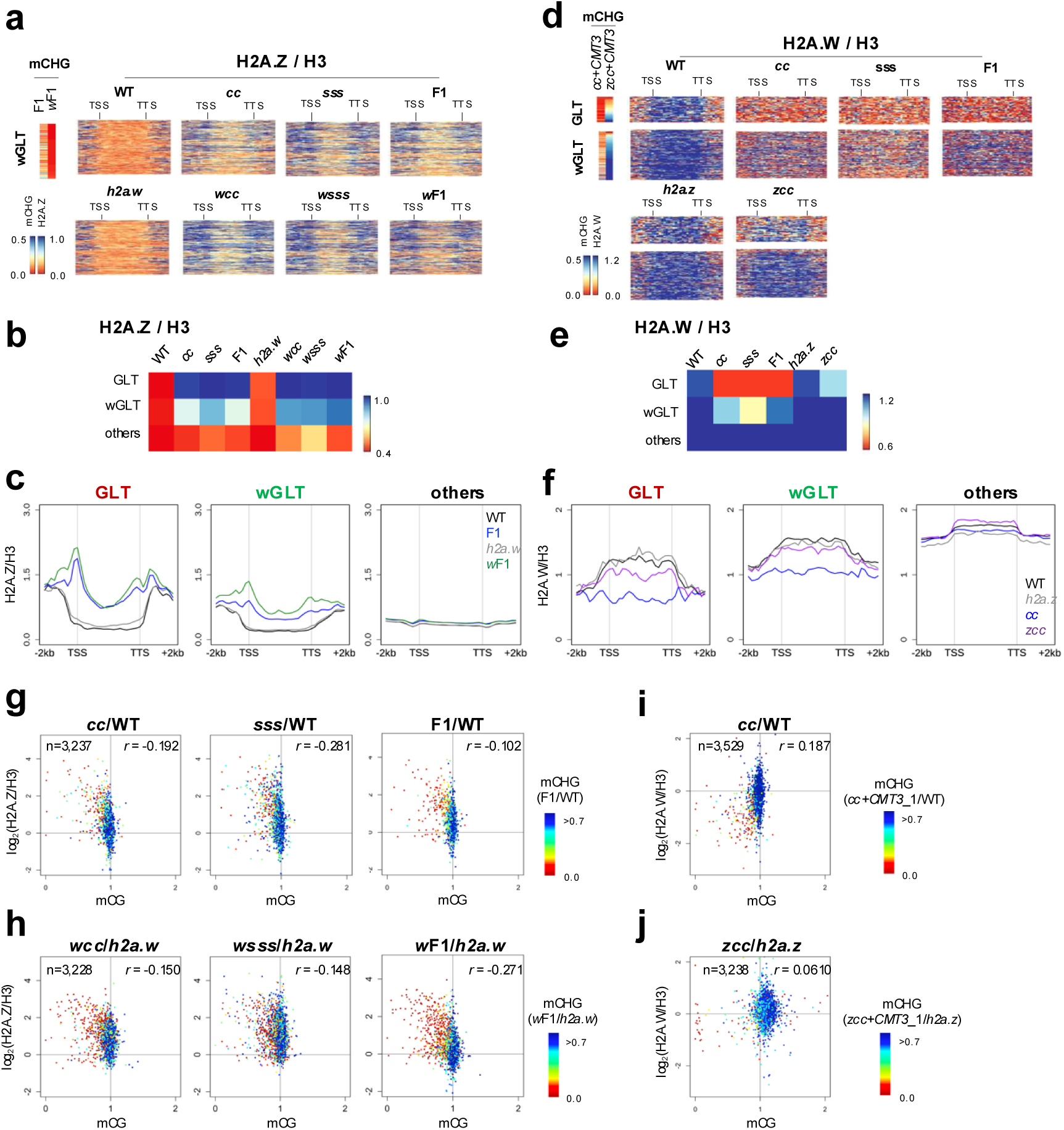
Mutual localization of H2A.W and H2A.Z in the absence of CMTs/SUVHs. **a** Heatmaps of H2A.Z over H3 within wGLTs and their flanking regions (2 kb) for WT, *h2a.w*, *cc*, *wcc*, *sss*, *wsss*, F1 and *w*F1 (n=208). The mean RPKM value for each segment was normalized to that of H3. TEs with a length > 1000 and mCHG in both WT and *h2a.w* > 0.1 were analyzed. TEs are sorted according to the mCHG levels in *w*F1. **b** Heatmaps showing the mean H2A.Z levels over H3 within TEs in the indicated genotypes (GLTs: n=73, wGLTs: n=232, other: n=3,598). **c** Metaplots of H2A.Z over H3 (GLTs: n=73, wGLTs: n=232, other: n=3,598). **d** Heatmaps showing H2A.W over H3 within and flanking regions (2 kb) of GLTs and wGLTs in WT, *cc, sss,* F1, *h2a.z* and *zcc* (GLTs: n=49, wGLTs: n=139). The mean RPKM value for each segment was normalized to that of H3. TEs with length > 1000 and mCHG in both WT and *h2a.z* > 0.1 were analyzed. TEs within each group are sorted according to the mCHG levels in *zcc*+*CMT3*_1. **e** Heatmaps showing the mean H2A.W levels over H3 within TEs in the indicated genotypes (GLTs: n=67, wGLTs: n=219, other: n=3,428). **f** Metaplots showing H2A.W over H3 (GLTs: n=73, wGLTs: n=232, other: n=3,598). **g,h** Comparison between mCG and H2A.Z/H3 in *cc*, *sss* and F1 relative to WT (**g**) and in *wcc*, *wsss* and *w*F1 relative to *h2a.w* (**h**). To avoid division by values near zero, TEs with mCHG, mCG, and H2A.Z/H3 (all >0.1) in WT (n=3,237) (**g**) or in *h2a.w* (n=3,228) (**h**) are shown, respectively. Each TE is colored according to the mCHG recovery level in F1 (F1/WT) (**g**) or in *w*F1 (*w*F1*/h2a.w*)(**h**). Spearman correlation coefficient was calculated and shown. **i,j** Comparison between mCG and H2A.W/H3 levels in *cc* relative to WT (**i**) and in *zcc* relative to *h2a.z* (**j**). To avoid division by values near zero, TEs with mCHG, mCG, and H2A.Z/H3 (all >0.1) in WT (n=3,529) (**i**) or in *h2a.z* (**j**) (n=3,238) are shown, respectively. Each TE is colored according to the mCHG recovery level in *cc*+*CMT3*_1 (*cc*+*CMT3*_1/WT) (**i**) or in *zcc*+*CMT3*_1 (*zcc*+*CMT3*_1*/h2a.z*) (**j**). Spearman correlation coefficient was calculated and shown.

Next, we examined the impact of *h2a.z* mutation on H2A.W localization. In *h2a.z*, H2A.W distribution was not significantly altered. In *cc*, *sss* and F1, H2A.W was lost at GLTs, as well as reduced in wGLTs (Fig. 3d-f, S6c,d). Intriguingly, this loss of H2A.W in *cc* was strongly suppressed by *h2a.z* mutation. In *zcc*, H2A.W enrichments significantly increased and became comparable to that in WT (Fig. 3d-f, S6c,d), suggesting that ectopic H2A.Z could decrease H2A.W in *cc*, and presumably as well in *sss* and F1. Furthermore, CG hypomethylation was also suppressed in *zcc* (Fig. 3i,j). Considering that *h2a.z* mutation also suppressed the failure of mCH recovery in GLTs and wGLTs (Fig. 2), these results suggest that H2A.Z localized ectopically in *CMTs/SUVHs* mutants can antagonize H2A.W and mCG only in GLTs and wGLTs, which could thereby inhibit mCH establishment in F1 and *w*F1.

Taken together, our results suggest that H2A.W and H2A.Z antagonistically affect each other for their localization at GLTs and wGLTs, particularly in the absence of mCH/H3K9me. Once this change has been induced, it is sustained even after complementation of *CMTs/SUVHs*, which would affect the dynamics of mCH/H3K9me establishment in wGLTs, and GLTs.

### Chromosomal environment and TE family influence the dynamics of mCH recovery

The results above suggest that mutually antagonistic H2A.Z and H2A.W affect mCH recovery dynamics, in GLTs and wGLTs. Conversely, they barely affected the other TEs in neither their mCH recovery nor the localization of the other variant. We hypothesized that GLTs and wGLTs have distinct properties, which would account for their unique dynamics of enrichment pattern in H2A variants and mCH recovery. One possibility would be chromosomal positions, as GLTs are predominantly distributed in chromosomal arms, where TE density is low compared to the pericentromeric regions^25^ (Fig. 4a, S7a). Furthermore, wGLTs are also enriched in the arms and around the edges of pericentromeric heterochromatin (Fig. 4a, S7a). GLTs are the farthest from centromeres, followed by wGLTs, and then the other TEs (Fig. 4b) which correlates positively with TE density and negatively with gene density (Fig. 4c). Thus, features associated with chromosomal positions may define GLTs and wGLTs.

**Fig. 4.**
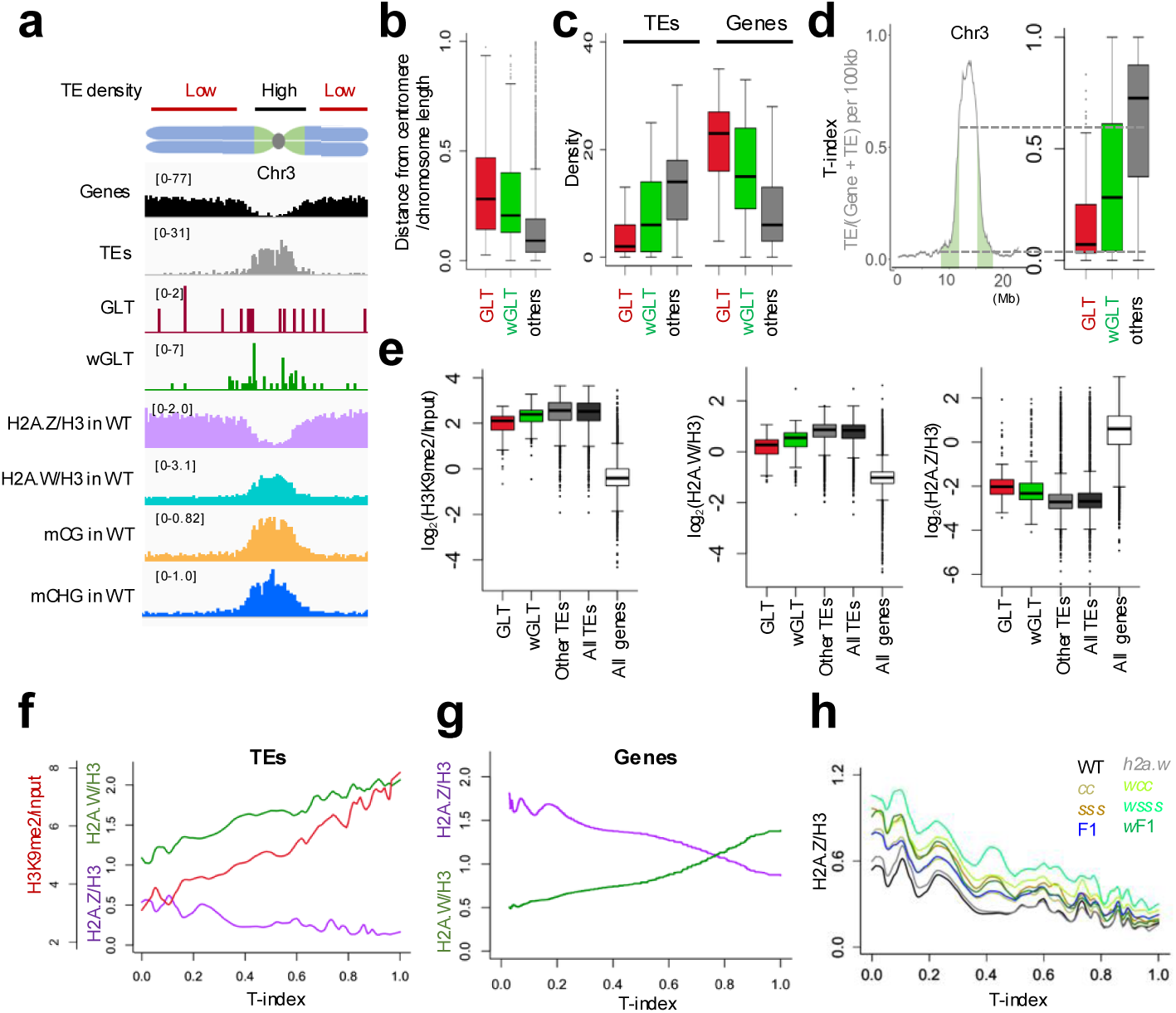
Chromosomal environment around TEs affects the dynamics of mCH recovery. **a** Genomic distributions of GLTs and wGLTs on Chr3. The numbers of normal protein-coding genes (black), TEs (gray), H2A.Z/H3 (purple), H2A.W/H3 (green), mCG (orange), and mCHG (blue) all in WT are shown in a 100-kb bin. **b** Distance from the centromere divided by the respective chromosome length in GLTs (red, n=73), wGLTs (green, n=232) and other TEs with mCHG in WT (gray, n=3,598). **c** The numbers of TEs (left) and normal protein-coding genes (right) around GLTs, wGLTs, and the other TEs (for 50 kb each side for a total 100 kb). **d** T-index (i.e. the number of TEs divided by the sum of TEs and genes in a 100-kb bin) over Chr3 (left). Box plots of T-index for GLTs, wGLTs, and the other TEs (right). The dashed lines indicate the first and third quartiles of the T-index for wGLTs. The green areas in the left figure indicate the regions with T-index between the first and third quartiles of wGLTs. **e** Enrichment of H3K9me2/input (left), H2A.W/H3 (middle) and H2A.Z/H3 (right) in WT for GLTs, wGLTs, the other TEs, all TEs with mCHG in WT > 0.1 (dark gray), and all genes without mCHG in WT (< 0.05)(white). **f** LOESS curves for H2A.Z/H3 (purple), H2A.W/H3 (green) and H3K9me2/input (red) levels of TEs in WT against T-index. The ChIP-seq data for H3K9me2 are from GSE148753. TEs with mCHG > 0.1 in WT were analyzed (n=3,598). **g** LOESS curves for H2A.Z/H3 (purple) and H2A.W/H3 (green) levels of protein-coding genes in WT (WT mCHG < 0.05, n=26,749) against T-index. **h** LOESS curves for H2A.Z/H3 of TEs against T-index. TEs with mCHG > 0.1 in both WT and *h2a.w* (n=3,511) were analyzed.

Chromosomal characteristics can vary among chromosomes, such as the length of chromosomes and the existence of pericentromeric knob in Chr 4^37^. To better represent ‘the degree of heterochromatin’, we introduced an index of TE density (T-index), which is calculated as numbers of TEs/(TEs + genes), in a 100-kb window (Fig. 4d, S7b left). As expected, T-index is strongly correlated with heterochromatic mark H3K9me2 (Table S1), as well as effectively capturing the pericentromeric knob feature around coordinates 2 Mb in Chr 4. T-index exhibits a steep increase and decrease around pericentromeric heterochromatin (hereafter we call these steep regions ‘boundaries’)(Fig. S7b, left). Next, T-index around each TE (50-kb on each side) was calculated, and then compared among GLTs, wGLTs and other TEs to the genomic T-index patterns. T-index for GLTs is low as in chromosomal arms. T-index for wGLTs overlaps the steeply changing boundaries. The other TEs show high T-index, as most of them are in pericentromeric regions (Fig. 4d, S7b right). These results reflect that GLTs predominantly localize in euchromatic arms, wGLTs around the boundaries, and the other TEs in pericentromeric heterochromatin.

Interestingly, even in WT, GLTs and wGLTs already show some euchromatic features; H3K9me2 and H2A.W are the lowest in GLTs, moderately lower in wGLTs than the other TEs, while the trend is anti-correlated in H2A.Z (Fig. 4e). The relationship between the heterochromatic features and T-index is seen beyond GLTs and wGLTs. LOESS-fitting curve shows that both H2A.W and H3K9me2 are positively correlated, while H2A.Z is negatively correlated with T-index (Fig. 4f). A similar trend was observed when GLTs and wGLTs were excluded from the analysis (Fig. S7c). The epigenetic features in protein-coding genes were also similarly affected by T-index, while the overall levels differed considerably between genes and TEs (Fig. 4e,g). These results suggest that, in both TEs and genes, epigenetic features can vary depending on the “chromosomal environments”, including surrounding chromatin structure (heterochromatin or euchromatin) and proximity to other genes or TEs. The stronger gene-like features in TEs with lower T-index were further enhanced in *cc*, *sss* and F1 (Fig. 4h, S7d). However, they largely reflect the changes in GLTs and wGLTs as the changes diminished greatly when GLTs and wGLTs were excluded (Fig. S7e,f), indicating that the acquisition of euchromatin-like properties in mCH mutants occurs particularly in GLTs and wGLTs.

TE family could be another possibility that may also account for the dependence on H2A variants for mCH recovery in GLTs and wGLTs (Fig. S8a). GLTs are enriched with *COPIA* and *LINE* retrotransposon families^25^. wGLTs are also enriched with *COPIAs* and *LINEs* but also with DNA transposon families. In both GLTs and wGLTs, *Gypsy* retrotransposons are underrepresented. The dynamics of mCH recovery and its dependence on H2A patterns varied significantly among TE families (Fig. S8b). *COPIAs* showed the least and *LINEs* the second least efficient mCH recovery, especially when H2A.W is absent, which is mostly suppressed when H2A.Z is absent (Fig. S8c).

*COPIAs* and *LINEs* tend to be distributed in chromosome arms, while DNA transposons are in pericentromeric heterochromatin, and *Gypsys* are in deep pericentromere^8,37,38^. By considering the distribution bias, the data was further dissected by T-index. Still, mCH recovery in *COPIAs* and *LINEs* was strongly dependent on both T-index and H2A.W. Even among TE copies with similar sequences, mCH recovery tended to be associated with T-index (Fig. S8d,e). Furthermore, GLTs and wGLTs exhibited a higher degree of enrichment with *COPIAs* and *LINEs* than TEs with low T-index (Fig. S8f). Taken together, the results suggest that both euchromatic environments represented by low T-index and TE families, *COPIAs* and *LINEs,* can best explain the specific displacement of H2A variants, which is associated with the failure of mCH recovery in GLTs and wGLTs.

### The restoration dynamics of heterochromatin states depend on T-index

The results above illuminate that the enrichment levels of H2A.W and H2A.Z (H2A patterns) is important for mCH establishment in TEs on chromosome arms and boundaries. On the contrary, pericentromeric TEs sustained TE-like H2A patterns (i.e. high H2A.W and/or low H2A.Z) even when both mCH and the other variant are absent. For example, the simultaneous loss of H2A.W and CMTs/SUVHs does not increase H2A.Z enrichment in pericentromeric TEs, obscuring the relationship of H2A variants and mCH recovery in pericentromeric regions. We wondered, if the replacement of H2A.W with H2A.Z occurs more genome-wide including pericentromeric heterochromatin, how does it affect the dynamics of mCH recovery? In order to address the question, we utilized the mCH/H3K9me reconstitution system in *met1-1*, a weak allele of mCG methyltransferase; the parental mutants *met1-1 cmt2 cmt3* mutant (*mcc*) and *met1-1 suvh4 suvh5 suvh6* mutant (*msss*), and their progeny *m*F1^39^. This is because our previous study shows that mCG decrease by *met1-1* mutation significantly attenuates mCH recovery more genome-wide including pericentromeric TEs^39^, and because mCG is known to antagonize H2A.Z^6,27^. We performed H2A.Z and H2A.W ChIP-seq, as well as re-analyzing WGBS data (GSE181896^39^). Overall, mCG decreased significantly in *met1-1* background (i.e. in *mcc*, *msss*, and *m*F1; Fig. 5a). This caused some increase in H2A.Z (Fig. 5b) as shown previously^27^, a slight decrease in H2A.W (Fig. 5c, S9c), and attenuation of mCH recovery (Fig. 5d), implying that the localization of H2A variants changed in a wide range of TEs, including those in pericentromeric heterochromatin.

**Fig. 5.**
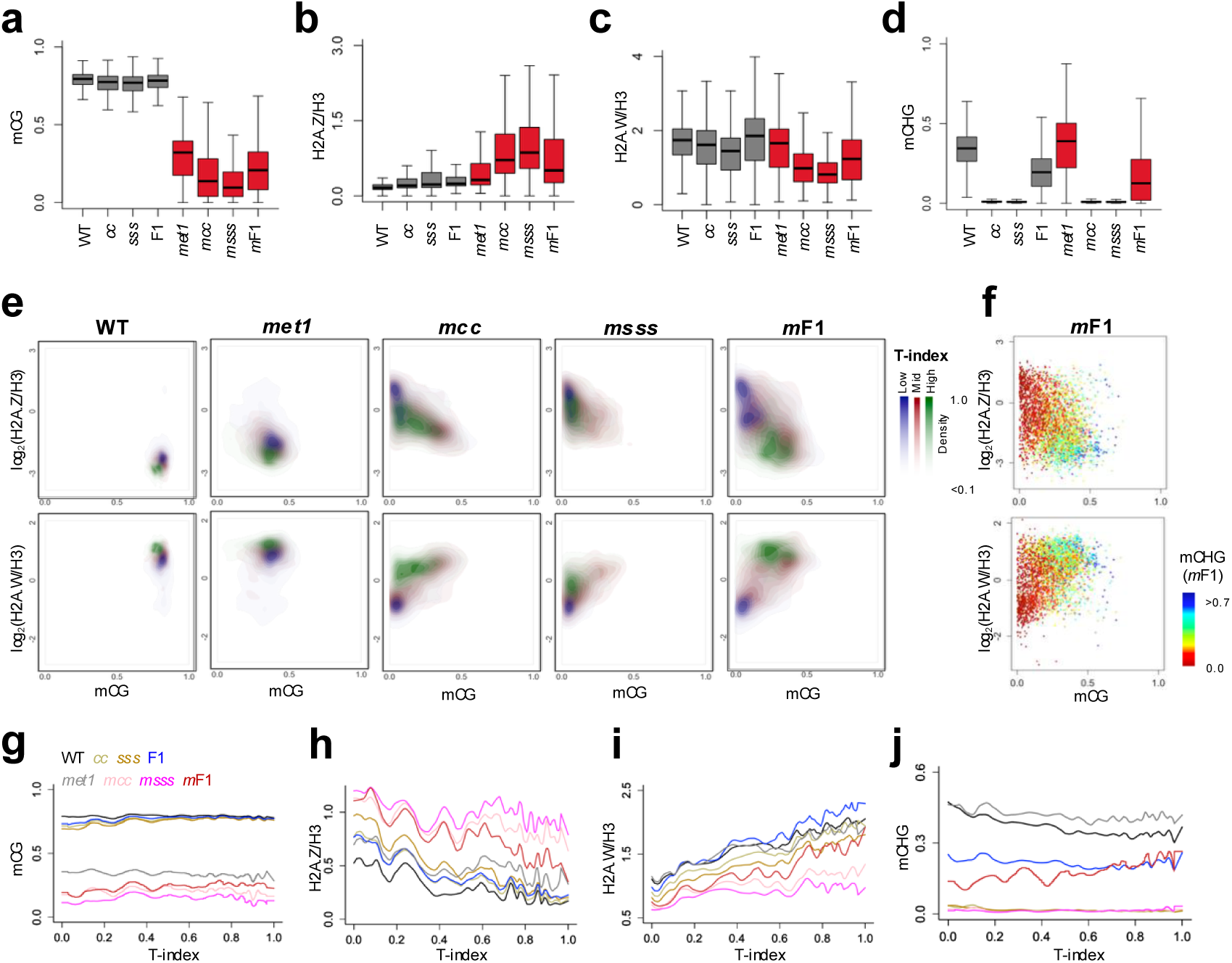
The restoration dynamics of heterochromatic marks are affected by T-index. **a-d** The levels of mCG (**a**), H2A.Z/H3 (**b**), H2A.W/H3 (**c**) and mCHG (**d**) in TEs in the indicated genotypes. **e** Density plots for the enrichments of H2A.Z/H3 (upper) and H2A.W/H3 (lower) against mCG levels, comparing among three groups of TEs (blue for low T-index (< 0.33); red for middle T-index (0.33 ≤ T-index < 0.67), green for high T-index ≥ 0.67). **f** Scatter plots of H2A.Z/H3 (upper) and H2A.W/H3 (lower) against mCG in each TE in *m*F1. Each TE is colored according to the mCHG levels in *m*F1. TEs with mCHG in both WT and *met1-1* > 0.1 (n=3,212) were analyzed. **g-j** LOESS curves for the levels of mCG, H2A.Z/H3, H2A.W/H3 and mCHG in TEs against T-index. TEs with mCHG in both WT and *met1-1* > 0.1 (n=3,212) were analyzed. DNA methylation data (GSE181896) were re-analyzed.

To investigate the relationship among chromosomal environments, H2A patterns, and mCG/mCH dynamics, TEs were categorized into three groups based on T-index. Then, ChIP-seq data of H2A.Z and H2A.W was compared against mCG levels as density plots (Fig 5e, S9a), scatter plots (Fig. 5f), as well as LOESS plots (Fig. 5g–j). As shown previously^25^, in the original system (i.e. *cc*, *sss*, and F1), a decrease in both mCG and H2A.W, and an increase in H2A.Z was observed in a subset of TEs with low T-index (shallow blue spreading; most of them reflect GLTs; Fig. S9a), which is linked to the absence of mCH recovery (Fig. S9b). In *met1-*1, mCG decreased by half on average, which was commonly observed irrespective of T-index (Fig. 5a,e,g). The *met1-1* mutation affected distribution of H2A variants slightly (Fig. 5b,c,e), with the exceptions for some TEs with substantial mCG and mCH loss (Fig. S9c). Interestingly, in *mcc* and *msss,* mCG further decreased (Fig. 5a,e,g), and this was associated with a significant increase in H2A.Z and a decrease in H2A.W, exhibiting a clear anti-correlation (Fig. 5b,c,e, S9d). Of particular significance was that displacement of H2A.W with H2A.Z occurred across the entire chromosomes, resulting in the majority of TEs acquiring euchromatic epigenetic states, irrespective of T-index (Fig. 5e,h,i). This allowed us to evaluate the impacts of chromosomal environments and H2A variants on mCH recovery, including pericentromeric heterochromatin.

Unexpectedly, although H2A patterns in the parents were similarly disturbed among three T-index groups, mCH recovery in *m*F1 was strikingly influenced by T-index. mCH recovered substantially in TEs with high T-index, but scarcely in TEs with low T-index (Fig. 5f,j, Fig. S9e). More surprisingly, other chromatin properties took the same behavior. TEs with high T-index reverted to their original epigenetic states of heterochromatin, losing H2A.Z and reacquiring H2A.W/mCG, with levels in *m*F1 becoming comparable with its control, *met1-1* (Fig. 5e-green in the right panels, h,i, S9f). On the contrary, TEs with low T-index remained in euchromatic states with high H2A.Z and low H2A.W/mCG in *m*F1, which is associated with the absence of mCH recovery (Fig. 5e-blue in the right panels, h,i, S9f). This trend is consistently seen when GLTs/wGLTs were excluded from the analysis (Fig. S9g). These results suggest that, for the recovery of heterochromatic features, local H2A patterns could be more important in TEs on chromosome arms than in pericentromeric regions. In other words, reacquisition of *CMTs/SUVHs* in *m*F1 triggered the autonomous and synchronous recovery in all the examined heterochromatic features in TE-dense pericentromeric regions but not in TE-poor and gene-dense chromosome arms, suggesting that chromosomal environments including TE density could be pivotal in accelerating the establishment of heterochromatin.

### mCH/H3K9me reconstitution in *h2a.z* can induce ectopic heterochromatin in protein-coding genes

The above evidence strongly indicates that ectopic H2A.Z suppresses mCH establishment in TEs, particularly in TE-poor euchromatin in chromosome arms. Given that H2A.Z predominantly localizes in genes in euchromatin, we asked whether H2A.Z in genes may also play a role in preventing mCH. Indeed, ectopic mCHG was clearly evident in *zcc*+*CMT3*s in hundreds of genes unmethylated in WT (mCHG in WT <0.05), despite only a limited number in *cc*+*CMT3*s (Fig. 6a, S10a,b). Thus, H2A.Z apparently prevents *de novo* mCH accumulation in genes. The enrichment levels of H2A.Z was comparable in mCH-gained genes to those in the other genes without mCHG in WT (Fig. S10c). Interestingly, mCH-gained genes did not significantly overlap among different *z*T1 plants (Fig. 6b), indicating that its occurrence is stochastic. Ectopic mCG was also observed not only in *z*T1s, but also in the parents, *zcc* (Fig. S10d), suggesting that ectopic mCG precedes ectopic mCHG in the progenies, *z*T1s. In fact, ectopic mCG was well associated with ectopic mCHG in the same individual but not in the other *z*T1s (Fig. 6c), which is consistent with our previous report that mCG is required for mCH establishment by CMTs/SUVHs^39^.

**Fig. 6.**
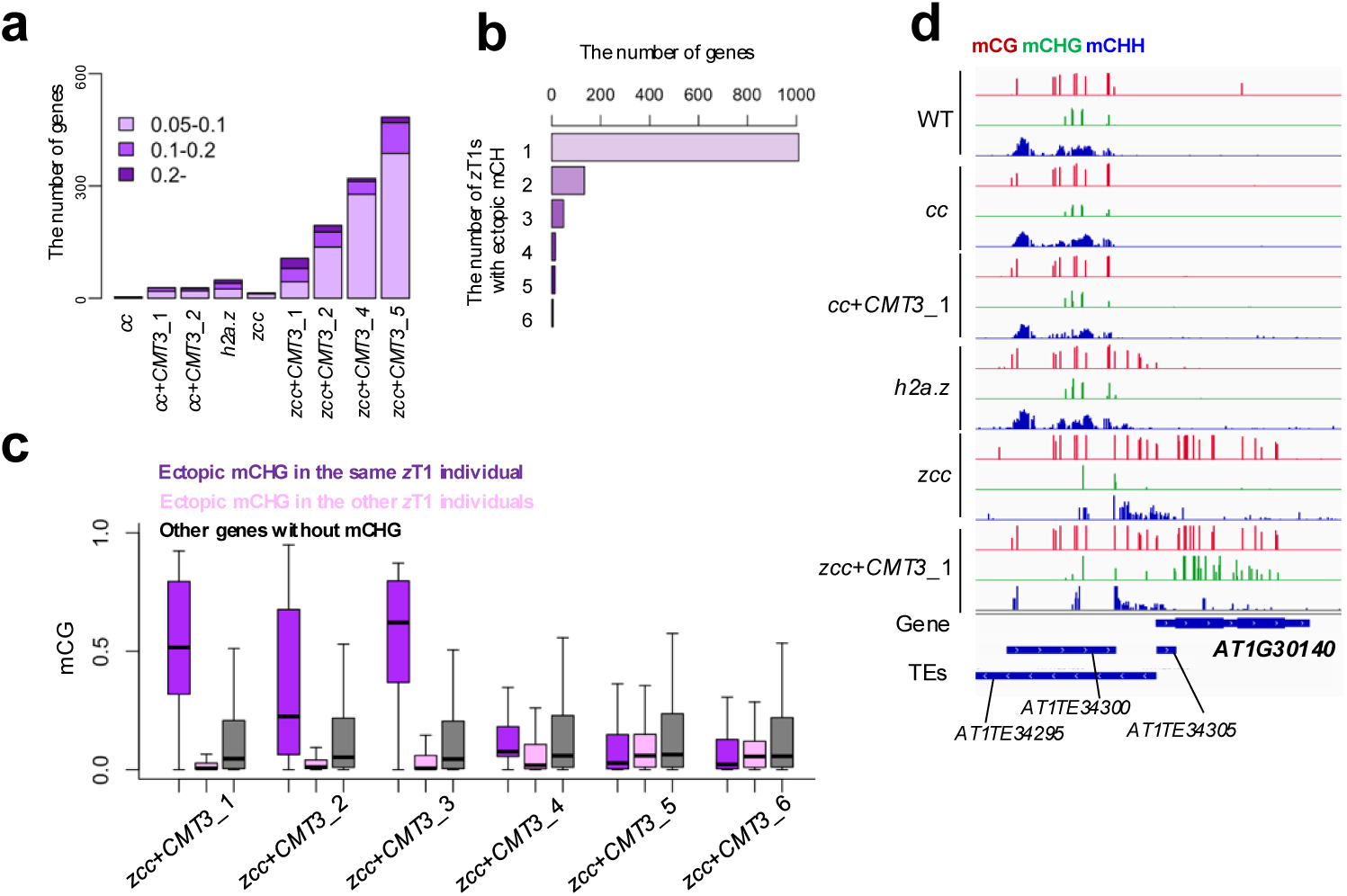
mCH/H3K9me reconstitution in *h2a.z* induces ectopic heterochromatin in protein-coding genes. **a** The numbers of genes with ectopic mCHG in the indicated plants. The genes with mCHG change (0.05-0.1, 0.1-0.2 and above 0.2) compared to WT were counted among the protein-coding genes without mCHG in WT (WT mCHG< 0.05). **b** Stochasticity for the ectopic mCHG in protein-coding genes. For each mCHG-gained gene in any individual *z*T1, the number of *z*T1 plants showing higher mCHG than WT (difference >0.05) were counted and shown. **c** mCG levels of genes with ectopic mCHG (difference >0.05) in the same *z*T1 individual compared with the genes with ectopic mCHG in the other *z*T1 individuals, and with other genes without ectopic mCHG in any *z*T1 individual. Outliers are excluded. **d** Genome browser view of mCG (red), mCHG (green) and mCHH (blue) in the indicated plants. Blue bars represent genes and TEs with TE fragments. The region Chr1/10,597,400-10,600,000 is shown.

Despite negligible levels of association to T-index (Fig. S10e), mCH-gained genes tended to localize in the vicinity of TEs (Fig. S10f). For instance, *AT1G30140* is located close to TEs with high DNA methylation (Fig. 6d). This gene is hypomethylated in WT, *cc* and the *cc*+*CMT3*. In *h2a.z*, mCG is slightly increased around the promoter. In *zcc*, mCG significantly increased towards the coding region of the gene, which is likely to be propagated from nearby TEs. Finally, in the *zcc*+*CMT3*, ectopic mCH was observed within the gene where mCG was present in the parent. Similar to this example, mCH-gained genes are frequently found in close proximity to TEs with high mCG and mCHG (Fig. S10g). The results suggest that the surrounding heterochromatic epigenetic state could affect nearby genes, increasing the chance of ectopic mCH accumulation, and that H2A.Z plays a pivotal role in preventing heterochromatin spreading.

## Discussion

In many eukaryotes, chromosomes are composed of distinct territories, euchromatin and heterochromatin. How this differential pattern is established in plants remains obscure, as RdDM predominantly functions in euchromatin. We previously proposed the RdDM-independent heterochromatin establishment mechanism, which precisely targets TEs for *de novo* mCH/H3K9me^25,26,39^. In this study, we demonstrate that mutually exclusive histone variants, H2A.W and H2A.Z, play a role in determining the targets of the *de novo* mCH/H3K9me mechanism in a subset of TEs; H2A.W promotes mCH establishment, while H2A.Z suppresses it. Interestingly, once chromatin becomes euchromatic (i.e. mCG/mCH/H2A.W is lost and H2A.Z is gained), heterochromatin re-formation is severely defected in TEs in chromosome arms, whereas heterochromatin is autonomously re-formed less dependently on H2A variants in TE-dense pericentromere. Moreover, H2A.Z in genes prevents heterochromatin spreading from proximal TEs. These results suggest the importance of histone H2A variants in regulating heterochromatin formation in euchromatin dominated regions of the genome (Fig. 7).

**Fig. 7.**
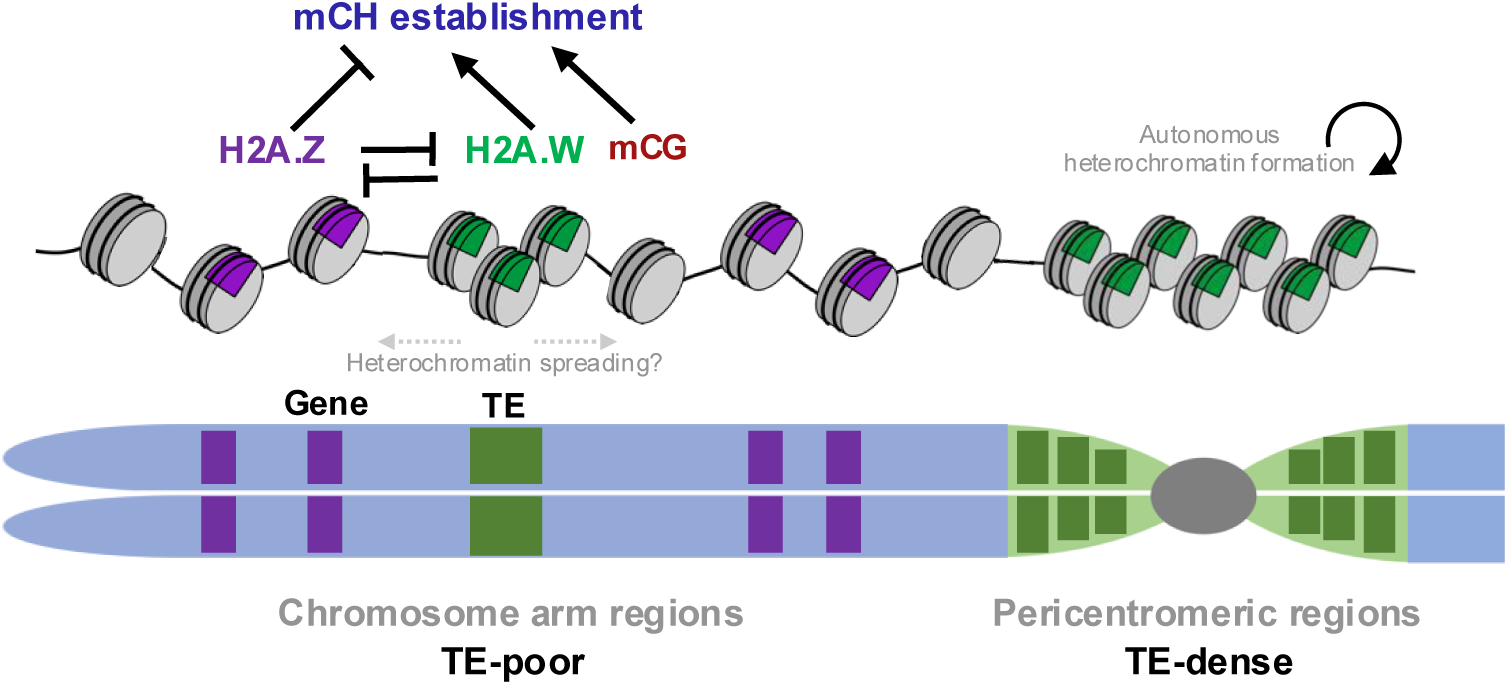
Model of specific heterochromatin formation at TEs in Arabidopsis. Heterochromatin is established specifically at TEs but not at genes. In chromosomal arm regions (TE-poor), H2A.W and H2A.Z, which antagonize each other, can promote and inhibit mCH establishment, respectively, at TEs. H2A.Z at genes can also prevent heterochromatin spreading into genes. Once chromatin becomes euchromatic genome-wide (i.e. with low mCG/mCH/H2A.W and high H2A.Z in *mcc* and *msss*), heterochromatin formation is severely defected in TEs in chromosome arms, while heterochromatin formation can occur autonomously in TE-dense pericentromeric regions. Thus, histone H2A variants play a pivotal role in regulating heterochromatin formation in euchromatin dominated regions of the genome.

Recent studies have shown that a subset of DDM1 functions in remodeling H2A variants by depositing H2A.W and possibly removing H2A.Z, although it is still unknown how DDM1 is recruited only to heterochromatic TEs. Despite the strong impacts shown in this study, how H2A variants contribute to *de novo* heterochromatin formation remains unclear. As either mutation alone rarely affects mCH distribution, these variants do not seem to directly regulate the activity of CMTs/SUVHs^28,30,31^. H2A.Z is known to be deposited at a proximity of nucleosome-free regions, and to reduce nucleosome stability, while H2A.W has the opposite effect^34,40,41^. Our results suggest that, when mCH is absent, ectopic H2A.Z can decrease H2A.W/mCG in hundreds of TEs in chromosomal arms, which could prevent the subsequent mCH establishment. In domains dominated by euchromatin, H2A.Z in genes may also prevent heterochromatin spreading into protein-coding genes from proximal TEs. Heterochromatin spreading can be deleterious, but has been reported in many eukaryotes^24,42–44^. H2A.Z could prevent such spreading into genes by modulating nucleosome-free regions, which is supported by the reports for a role of H2A.Z in preventing *de novo* DNA methylation in zebrafish embryos^45^, and as the boundary against heterochromatin in yeast^46^. In contrast to the local effects of H2A.Z, H2A.W, which shares the KSPK motif with mammalian macroH2A^41,47,48^, is more likely to regulate constitutive heterochromatin at genomic level; higher-order chromatin compaction^30,31^, chromatin accessibility together with H1^31^, and inhibition of mitotic crossover in heterochromatin^49^.

The distribution of TEs and genes, or heterochromatin and euchromatin, differs among species. In certain species including fruit fly, fission yeast, and Arabidopsis, heterochromatin is clustered at some restricted regions of chromosomes^24,37,43^. In species with larger genome size, including maize, mouse and human, TEs tend to be dispersed in chromosomes and are often found proximal to genes^50,51^, while their genomes are often functionally compartmented like TE-rich/poor regions. Important questions remain how H2A variants impact heterochromatin formation in other species including those with more abundant TEs in their genomes, and the extent to which the current findings apply.

## Methods

### Plant materials and growth conditions

Columbia-0 (Col-0) strain was used as the WT. The mutants *h2a.z* (*hta8 hta9 hta11*) and *cmt2 cmt3* (*cc*), *suvh4 suvh5 suvh6* (*sss*), and *met1-1* are kind gifts from Frederic Berger, Daniel Zilberman, Judith Bender, and Eric Richards^13,16,28,52^. The mutants *h2a.w-2* (*hta6 hta7 hta12*), *zcc*, *mcc*, *msss* and *m*F1 were described previously^25,31,39^. The mutants *wcc* and *wsss* were created in this study by crossing *h2a.w* and *cc* or *h2a.w* and *sss*, respectively. *w*F1was created by crossing *wcc* and *wsss. z*T1s were created by transforming *CMT3* and *CMT2* transgenes, respectively, into *zcc*^25^ (for *zcc+CMT3_1-3 and zcc+CMT2_1-3*) and into *ccz* (for *zcc+CMT3_4-6*)^25^. The plants were grown at 22°C under 16 h light and 8 h dark conditions for one to two weeks on MS agar medium containing 1% sucrose. They were then grown on soil under the same condition.

### Plasmid construction and plant transformation

For the transformation of *CMT3* and *CMT2* genes, the genomic regions from their promoter to just before stop codon were amplified with the following primers respectively: 5′-cgggccccccctcgaagcttgtatgattagatatgtgaa-3′ and 5′-ggatccacctccacccccgggtgcaagctcggaaggaagag-3′ for *CMT3*, 5′-cgggccccccctcgaagcttaacccgtagatatggcacaaag-3′ and 5′-ggatccacctccacccccgggatgaggaatggtttcttgaag-3′ for *CMT2*. The amplified fragments were cloned into HindIII-digested pGreenII-based vector with GFP by using NEBuilder HiFi DNA Assembly Master Mix (NEB). The plasmids were transformed into *Agrobacterium tumefaciens* using electroporation, and then into *cc*, *zcc* and *ccz* plants by the floral dip. The transgenic lines were selected on MS agar plates with Hygromycin and Carbenicillin. The selected plants were then grown on soil.

### RT-qPCR

Rosette leaves from three- to four-week-old individual plant were frozen with liquid nitrogen, ground, and total RNA was purified using TRIzol RNA Isolation Reagents (Thermo). RNA was treated with DNase I, reverse-transcribed into cDNA using SuperScript IV VILO Master Mix with ezDNase (Thermo). cDNA was amplified using KAPA SYBR FAST qPCR Master Mix with the following primers: 5′-gagacaaacgtgagaaacatgacc-3′ and 5′-cgctgacatcttcattgtcctc-3′ for *CMT3*, 5′-ccataagagaatctgcacggc-3′ and 5′-aagccataccaagcgaataacc-3′ for *CMT2* and 5′-gatctccaaggccgagtatgat-3′ and 5′-cccattcataaaaccccagc-3′ for *ACT2*. qPCR was performed using StepOnePlus Real-Time PCR System (Thermo). Two independent biological replicates were analyzed. The expression levels of *CMT3* and *CMT2* were normalized with the expression levels of *ACT2*. Bar plots in Figure S4f and S5k,l were created by ggplot2 (v3.3.2).

### Whole-genome bisulfite sequencing and data processing

Three- to four-week-old rosette leaves of one individual plant were frozen with liquid nitrogen, ground, and the genomic DNA was extracted using Nucleon PhytoPure (Cytiva). The genomic DNA was fragmented using S220 Focused Ultrasonicator (Covaris) and the size of 300-500bp was gel-extracted. The libraries were prepared using TruSeq DNA LT Sample Prep Kit (Illumina) and KAPA HyperPrep Kit (Kapa Biosystems) and then bisulfite conversion was performed using MethylCode Bisulfite Conversion Kit (Thermo). The libraries were amplified using KAPA HiFi HotStart Uracil ReadyMix (Kapa Biosystems) and purified using SPRIselect (Beckman Coulter). 150-bp paired-end sequencing was performed using the HiSeq X and Novaseq sequencer (Illumina) at Macrogen Japan Corp. Two independent biological replicates were analyzed. Raw sequence data were trimmed and quality-filtered using Trimmomatic version 0.33^53^. The trimmed and quality-filtered sequences were mapped to the reference genome (TAIR10^54^), deduplicated and analyzed to extract methylation data using Bismark version 0.10.1^55^. To calculate methylation levels in genes, TEs (TEs with protein-coding genes) and TEs including TE fragments, a custom Perl script was used^25,39^. CG, CHG and CHH methylation levels within each region were calculated as the number of methylated cytosines divided by the number of total cytosines (weighted methylation level)^56^. The regions with the number of total cytosines < 10 were excluded from the analysis. In addition, for mutants containing the *h2a.z* mutation, regions in which more than 80% of cytosines had fewer than 5 mapped reads were also excluded from the analysis. Plots were created using Rstudio version 1.3.959. Venn diagrams were created using BioVenn^57^. TEs with statistically lower or higher mCH in *w*F1 than in F1 or mCHG/mCHH in *zcc*+*CMT3*/*2* than in *cc*+*CMT3*/*2* shown in Figures 1b, 2g, S1b, S4g,h and S5g,n,o were extracted by using edgeR (v3.32.1)^58^ following Chen et al.^59^. For browser views, bedGraph files were created and visualized using IGV (v2.8.9)^60^.

### ChIP–seq and data processing

The antibodies used were rabbit anti-H2A.Z^25^, rabbit anti-H2A.W6^30^ and rabbit anti-H3 (ab1791; Abcam). For H2A.Z ChIP-seq, eChIP method^61^ was performed as described previously^62^. For H2A.W ChIP-seq, cross-linking and chromatin fragmentation were performed as previously^62^. Fragmented chromatin samples were incubated with anti-H2A.W or rabbit anti-H3 overnight at 4°C with rotation, followed by incubation with Dynabeads Protein G (invitrogen) for 2 hours at 4°C. These were washed once with low-salt ChIP buffer, twice with high-salt ChIP buffer, once with ChIP wash buffer and TE buffer, and eluted from beads by adding 150 μl of ChIP Elution buffer^61^ with incubation for 6-9 hours at 65°C. DNA was extracted from the eluted samples with Monarch PCR & DNA Cleanup Kit (NEB) and libraries were constructed with ThruPLEX DNA-Seq Kit (Takara). 150-bp paired-end sequencing was performed using the HiSeq X Ten and Novaseq sequencer (Illumina) at Macrogen Japan Corp. Two independent biological replicates were analyzed. Raw sequence data were trimmed and quality-filtered using Trimmomatic version 0.33^53^. The trimmed and quality-filtered sequences were mapped to the reference genome (TAIR10) using Bowtie (version 1.1.2) with the option -M 1–best^63^. BEDTools (version 2.26.0) was used to count the reads that overlapped with each gene, TE or TE including fragments^64^. Plots were created using Rstudio. To create metaplots and heatmaps around and within genes and TEs, 2kb upstream and downstream of genes and TEs were divided into 10 segments (200b) and TEs were divided into 20 segments. The mapped reads within each segment of TEs and the flanking regions were counted using bedtools and normalized as RPKM using Perl script. Metaplots were created with the averaged RPKM within each segment and plotted using R version 4.3.1. Heatmaps were created using gplots (v3.1.1).

### Phylogenetic analysis of *COPIA* copies

*COPIA* sequences were retrieved from TE annotation of TAIR9.0. Among 1800 annotated regions, copies with RT core domain region (ca.260 aa) were used. After manual alignment, sequences with more than 100bp gap were discarded. The remaining 202 sequences were used for calculation of genetic distances with p-distance. Evolutionary analyses were conducted by MEGA5^65^. TE pairs with a p-distance < 0.25 that are each other’s closest in terms of p-distance were selected. The pairs with mCHG in WT and *h2a.w* > 0.1 and with mCHG in WT and *h2a.z* > 0.1 were selected.

## Supporting information

Supplemental_Figs_Table_Reference

## Data availability

WGBS and ChIP-seq data generated in this study were deposited in the GEO with the accession number GSE303147. WGBS data and ChIP-seq data (H3K9me2) for WT, *cc*, *sss*, and F1 is available in the GEO with the accession number GSE148753^25^. WGBS data for *met1-1*, *mcc*, *msss* and *m*F1 is available in the GEO with the accession number GSE181896^39^. TAIR10^54^ (https://www.arabidopsis.org) was used as the Arabidopsis reference genome. Source data are provided with this paper.

## Code availability

The codes used in this study are available via contacting the corresponding author.

## Acknowledgements

We thank Daniel Zilberman, Judith Bender, and Eric Richards for sharing mutant strains. Computations were partially performed on the NIG supercomputer at ROIS National Institute of Genetics. This work was supported by grants from JST FOREST (JPMJFR224X to T.K.T), JST PRESTO (JPMJPR20K3 to A.O.), JST CRESTO (JPMJCR20S to A.O.), JSPS KAKENHI (17K15059, 19H05740, 21H04977, 22K06180 and 25H01299 to T.K.T. and 24H01351, 24K02077, 24K02002, 25K02258, 25H02544, and 25H01300 to A.O.), and JSPS Fellowship (23KJ0740 to S.O.).

## Author contributions

S.O., T.K., and T.K.T. designed the study. S.O., S.T, S.T, A.O., F.B., T.K., and T.K.T. performed the experiments. S.O., A.K., and T.K.T. analyzed the data. S.O. and T.K.T. wrote the paper incorporating comments from the other authors.

## References

1. Cosby, R. L., Chang, N.-C. & Feschotte, C. Host-transposon interactions: conflict, cooperation, and cooption. Genes Dev. 33, 1098–1116 (2019).

2. Kim, M. Y. & Zilberman, D. DNA methylation as a system of plant genomic immunity. Trends Plant Sci. 19, 320–326 (2014).

3. Cokus, S. J. et al. Shotgun bisulphite sequencing of the Arabidopsis genome reveals DNA methylation patterning. Nature 452, 215–219 (2008).

4. Lister, R. et al. Highly integrated single-base resolution maps of the epigenome in Arabidopsis. Cell 133, 523–536 (2008).

5. Zhang, X. et al. Genome-wide high-resolution mapping and functional analysis of DNA methylation in arabidopsis. Cell 126, 1189–1201 (2006).

6. Zemach, A., McDaniel, I. E., Silva, P. & Zilberman, D. Genome-wide evolutionary analysis of eukaryotic DNA methylation. Science 328, 916–919 (2010).

7. To, T. K., Saze, H. & Kakutani, T. DNA methylation within transcribed regions. Plant Physiol. 168, 1219–1225 (2015).

8. Underwood, C. J., Henderson, I. R. & Martienssen, R. A. Genetic and epigenetic variation of transposable elements in Arabidopsis. Curr. Opin. Plant Biol. 36, 135–141 (2017).

9. Zhang, H., Lang, Z. & Zhu, J.-K. Dynamics and function of DNA methylation in plants. Nat. Rev. Mol. Cell Biol. 19, 489–506 (2018).

10. Ma, R. et al. Targeting pericentric non-consecutive motifs for heterochromatin initiation. Nature 631, 678–685 (2024).

11. Jackson, J. P., Lindroth, A. M., Cao, X. & Jacobsen, S. E. Control of CpNpG DNA methylation by the KRYPTONITE histone H3 methyltransferase. Nature 416, 556–560 (2002).

12. Malagnac, F., Bartee, L. & Bender, J. An Arabidopsis SET domain protein required for maintenance but not establishment of DNA methylation. EMBO J. 21, 6842–6852 (2002).

13. Ebbs, M. L. & Bender, J. Locus-specific control of DNA methylation by the Arabidopsis SUVH5 histone methyltransferase. Plant Cell 18, 1166–1176 (2006).

14. Bartee, L., Malagnac, F. & Bender, J. Arabidopsis *cmt3* chromomethylase mutations block non-CG methylation and silencing of an endogenous gene. Genes Dev. 15, 1753– 1758 (2001).

15. Lindroth, A. M. et al. Requirement of CHROMOMETHYLASE3 for maintenance of CpXpG methylation. Science 292, 2077–2080 (2001).

16. Zemach, A. et al. The Arabidopsis nucleosome remodeler DDM1 allows DNA methyltransferases to access H1-containing heterochromatin. Cell 153, 193–205 (2013).

17. Stroud, H. et al. Non-CG methylation patterns shape the epigenetic landscape in Arabidopsis. Nat. Struct. Mol. Biol. 21, 64–72 (2014).

18. Johnson, L., Cao, X. & Jacobsen, S. Interplay between two epigenetic marks. DNA methylation and histone H3 lysine 9 methylation. Curr. Biol. 12, 1360–1367 (2002).

19. Du, J. et al. Dual binding of chromomethylase domains to H3K9me2-containing nucleosomes directs DNA methylation in plants. Cell 151, 167–180 (2012).

20. Finnegan, E. J. & Dennis, E. S. Isolation and identification by sequence homology of a putative cytosine methyltransferase from Arabidopsis thaliana. Nucleic Acids Res. 21, 2383–2388 (1993).

21. Finnegan, E. J., Peacock, W. J. & Dennis, E. S. Reduced DNA methylation in Arabidopsis thaliana results in abnormal plant development. Proc. Natl. Acad. Sci. U. S. A. 93, 8449–8454 (1996).

22. Matzke, M. A. & Mosher, R. A. RNA-directed DNA methylation: an epigenetic pathway of increasing complexity. Nat. Rev. Genet. 15, 394–408 (2014).

23. Cuerda-Gil, D. & Slotkin, R. K. Non-canonical RNA-directed DNA methylation. Nat. Plants 2, 16163 (2016).

24. Grewal, S. I. S. The molecular basis of heterochromatin assembly and epigenetic inheritance. Mol. Cell 83, 1767–1785 (2023).

25. To, T. K. et al. RNA interference-independent reprogramming of DNA methylation in Arabidopsis. Nat. Plants 6, 1455–1467 (2020).

26. To, T. K. & Kakutani, T. Crosstalk among pathways to generate DNA methylome. Curr. Opin. Plant Biol. 68, 102248 (2022).

27. Zilberman, D., Coleman-Derr, D., Ballinger, T. & Henikoff, S. Histone H2A.Z and DNA methylation are mutually antagonistic chromatin marks. Nature 456, 125–129 (2008).

28. Coleman-Derr, D. & Zilberman, D. Deposition of histone variant H2A.Z within gene bodies regulates responsive genes. PLoS Genet. 8, e1002988 (2012).

29. Jamge, B. et al. Histone variants shape chromatin states in Arabidopsis. eLife 12, (2023).

30. Yelagandula, R. et al. The histone variant H2A.W defines heterochromatin and promotes chromatin condensation in Arabidopsis. Cell 158, 98–109 (2014).

31. Bourguet, P. et al. The histone variant H2A.W and linker histone H1 co-regulate heterochromatin accessibility and DNA methylation. Nat. Commun. 12, 2683 (2021).

32. Osakabe, A. et al. The chromatin remodeler DDM1 prevents transposon mobility through deposition of histone variant H2A.W. Nat. Cell Biol. 23, 391–400 (2021).

33. Zhou, J. et al. DDM1-mediated R-loop resolution and H2A.Z exclusion facilitates heterochromatin formation in Arabidopsis. Sci. Adv. 9, eadg2699 (2023).

34. Osakabe, A. et al. Molecular and structural basis of the chromatin remodeling activity by Arabidopsis DDM1. Nat. Commun. 15, 5187 (2024).

35. Bourguet, P. et al. The histone variant H2A.W cooperates with chromatin modifications and linker histone H1 to maintain transcriptional silencing of transposons in Arabidopsis. Preprint at https://www.biorxiv.org/content/10.1101/2022.05.31.493688v1 (2022).

36. Wang, Y. et al. Structural and functional interrelationships of histone H2A with its variants H2A.Z and H2A.W in Arabidopsis. Structure 33, 1240–1249.e5 (2025).

37. Arabidopsis Genome Initiative. Analysis of the genome sequence of the flowering plant Arabidopsis thaliana. Nature 408, 796–815 (2000).

38. Quesneville, H. Twenty years of transposable element analysis in the Arabidopsis thaliana genome. Mob. DNA 11, 28 (2020).

39. To, T. K. et al. Local and global crosstalk among heterochromatin marks drives DNA methylome patterning in Arabidopsis. Nat. Commun. 13, 861 (2022).

40. Osakabe, A. et al. Histone H2A variants confer specific properties to nucleosomes and impact on chromatin accessibility. Nucleic Acids Res. 46, 7675–7685 (2018).

41. Talbert, P. B. & Henikoff, S. Histone variants at a glance. J. Cell Sci. 134, jcs244749 (2021).

42. Diez, C. M., Roessler, K. & Gaut, B. S. Epigenetics and plant genome evolution. Curr. Opin. Plant Biol. 18, 1–8 (2014).

43. Mteirek, R., Gueguen, N., Jensen, S., Brasset, E. & Vaury, C. Drosophila heterochromatin: structure and function. Curr. Opin. Insect Sci. 1, 19–24 (2014).

44. Nicetto, D. & Zaret, K. S. Role of H3K9me3 heterochromatin in cell identity establishment and maintenance. Curr. Opin. Genet. Dev. 55, 1–10 (2019).

45. Murphy, P. J., Wu, S. F., James, C. R., Wike, C. L. & Cairns, B. R. Placeholder nucleosomes underlie germline-to-embryo DNA methylation reprogramming. Cell 172, 993–1006.e13 (2018).

46. Meneghini, M. D., Wu, M. & Madhani, H. D. Conserved histone variant H2A.Z protects euchromatin from the ectopic spread of silent heterochromatin. Cell 112, 725–736 (2003).

47. Berger, F., Muegge, K. & Richards, E. J. Seminars in cell and development biology on histone variants remodelers of H2A variants associated with heterochromatin. Semin. Cell Dev. Biol. 135, 93–101 (2023).

48. Wong, L. H. & Tremethick, D. Multifunctional histone variants in genome function. Nat. Rev. Genet. 26, 82–104 (2025).

49. Son, N. et al. The histone variant H2A.W restricts heterochromatic crossovers in Arabidopsis. Proc. Natl. Acad. Sci. U. S. A. 122, e2413698122 (2025).

50. Lawson, H. A., Liang, Y. & Wang, T. Transposable elements in mammalian chromatin organization. Nat. Rev. Genet. 24, 712–723 (2023).

51. Schnable, P. S. et al. The B73 maize genome: complexity, diversity, and dynamics. Science 326, 1112–1115 (2009).

52. Kankel, M. W. et al. Arabidopsis MET1 cytosine methyltransferase mutants. Genetics 163, 1109–1122 (2003).

53. Bolger, A. M., Lohse, M. & Usadel, B. Trimmomatic: a flexible trimmer for Illumina sequence data. Bioinformatics 30, 2114–2120 (2014).

54. Lamesch, P. et al. The Arabidopsis Information Resource (TAIR): improved gene annotation and new tools. Nucleic Acids Res. 40, D1202–10 (2012).

55. Krueger, F. & Andrews, S. R. Bismark: a flexible aligner and methylation caller for Bisulfite-Seq applications. Bioinformatics 27, 1571–1572 (2011).

56. Schultz, M. D., Schmitz, R. J. & Ecker, J. R. ‘Leveling’ the playing field for analyses of single-base resolution DNA methylomes. Trends Genet. 28, 583–585 (2012).

57. Hulsen, T., de Vlieg, J. & Alkema, W. BioVenn - a web application for the comparison and visualization of biological lists using area-proportional Venn diagrams. BMC Genomics 9, 488 (2008).

58. Robinson, M. D., McCarthy, D. J. & Smyth, G. K. edgeR: a Bioconductor package for differential expression analysis of digital gene expression data. Bioinformatics 26, 139– 140 (2010).

59. Chen, Y., Pal, B., Visvader, J. E. & Smyth, G. K. Differential methylation analysis of reduced representation bisulfite sequencing experiments using edgeR. F1000Res. 6, 2055 (2017).

60. Robinson, J. T., et al. Integrative genomics viewer. Nat. Biotechnol. 29, 24–26 (2011).

61. Zhao, L. et al. Integrative analysis of reference epigenomes in 20 rice varieties. Nat. Commun. 11, 2658 (2020).

62. Mori, S. et al. Cotranscriptional demethylation induces global loss of H3K4me2 from active genes in Arabidopsis. EMBO J. 42, e113798 (2023).

63. Langmead, B., Trapnell, C., Pop, M. & Salzberg, S. L. Ultrafast and memory-efficient alignment of short DNA sequences to the human genome. Genome Biol. 10, R25 (2009).

64. Quinlan, A. R. & Hall, I. M. BEDTools: a flexible suite of utilities for comparing genomic features. Bioinformatics 26, 841–842 (2010).

65. Tamura, K. et al. MEGA5: molecular evolutionary genetics analysis using maximum likelihood, evolutionary distance, and maximum parsimony methods. Mol. Biol. Evol. 28, 2731–2739 (2011).

