## Supplemental_Figs_Table_Reference for "Antagonistic histone H2A variants and autonomous heterochromatin formation shape epigenomic patterns in Arabidopsis"

### Fig.S1

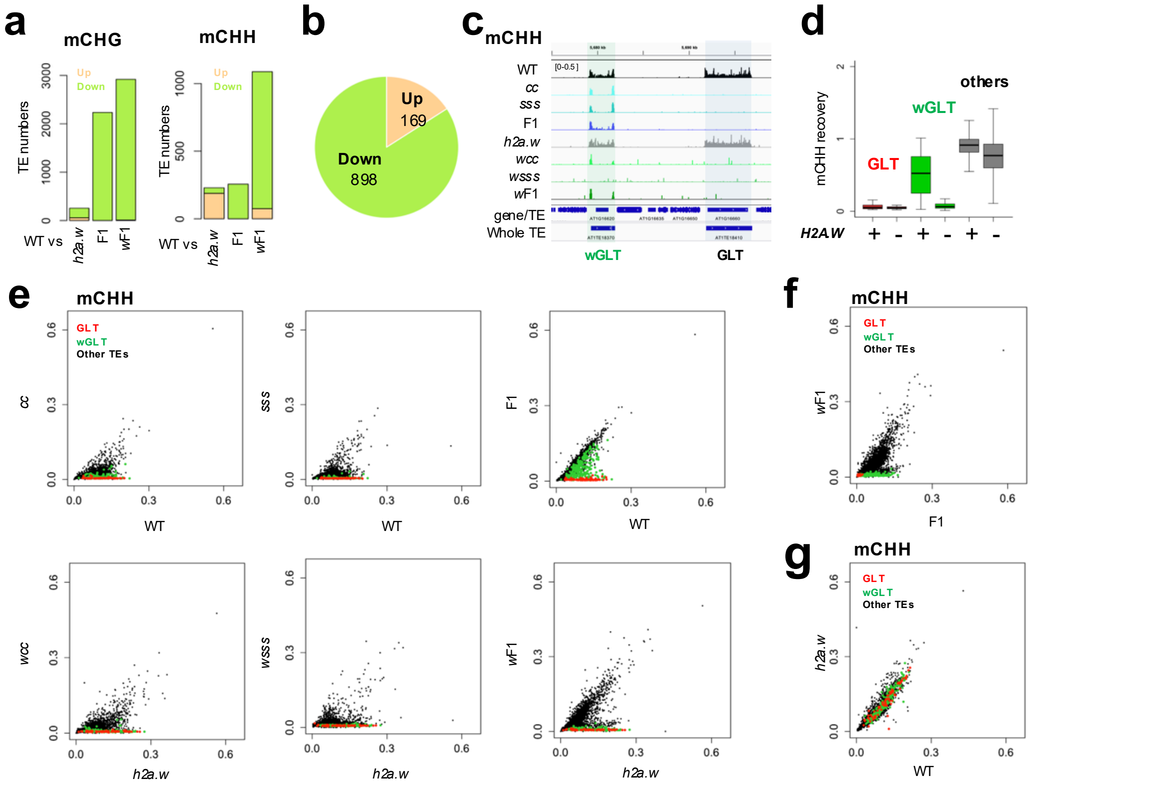


**_h2a.w_ mutation reduces the efficiency of mCHH recovery in hundreds of TEs**

**a** Bar plots for the number of TEs with significant changes in mCHG (left) or mCHH (right) in *h2a.w*, F1 and *w*F1 compared to WT. **b** Pie chart showing the number of TEs with statistically significant changes in mCHH in *w*F1 compared to the original F1. **c** Genome browser view of mCHH in the indicated plants as in the format in Fig. 1**c**. **d** Comparison of mCHH recovery in F1 and *w*F1. The recovery was calculated as F1/WT (H2A.W +) and *w*F1/*h2a.w* (H2A.W -), respectively. To avoid division by values near zero, TEs with mCHH (>0.03) in both WT and *h2a.w* were used (n = 3350). Outliers are not shown. The centerline and box edges represent quartiles and whiskers range 1.5 times of the interquartile from the box edges. **e** mCHH level of each TE in *cc* (left), *sss* (middle), and the F1 (right) compared to WT as well as *wcc* (left), *wsss* (middle), and the *w*F1 (right) compared to *h2a.w*. GLTs are indicated in red, wGLTs in green, and the other TEs in black. **f,g** Comparison of mCHH levels for each TE between the F1 and *w*F1 (**f**) and WT and *h2a.w* (**g**).

### Fig.S2

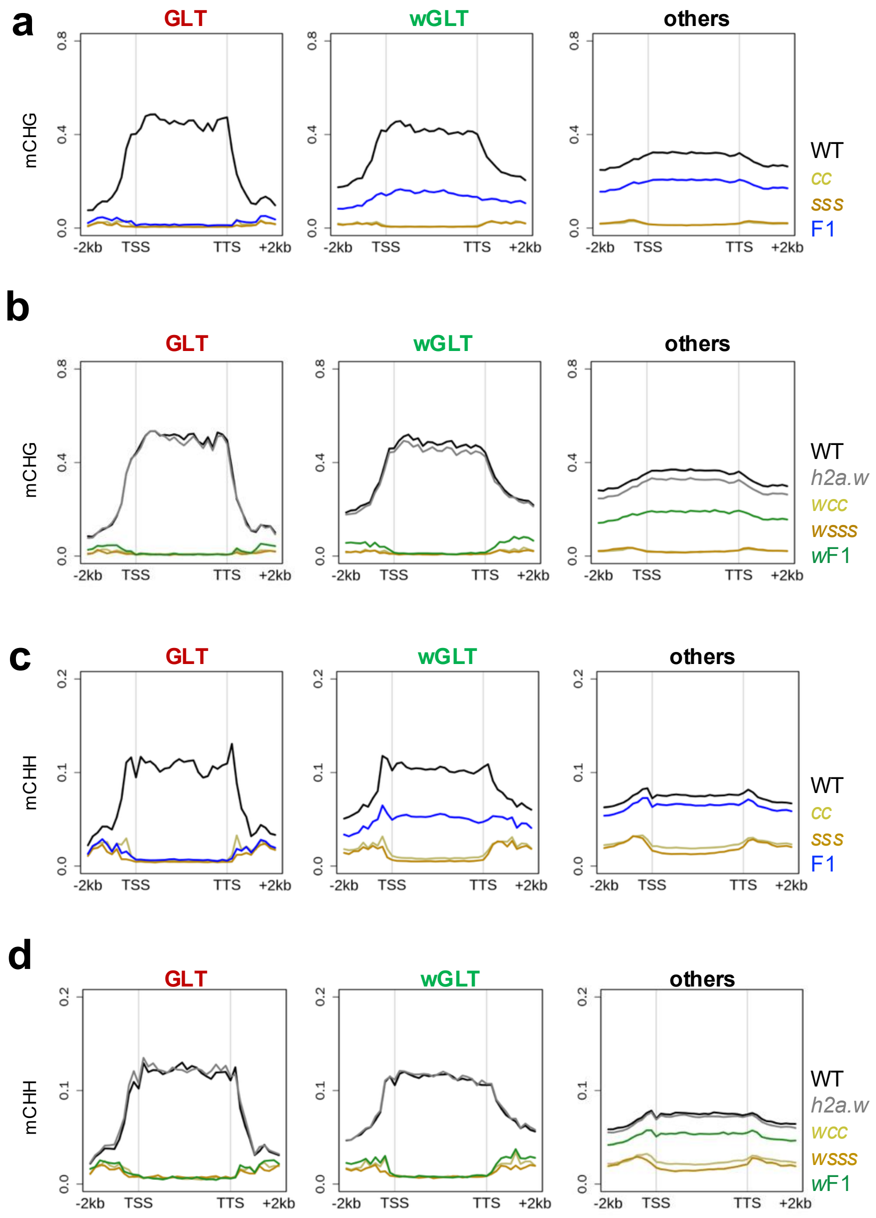


**Metaplots of mCH in GLTs, wGLTs and other TEs**
Metaplots of mCHG (**a**,**b**) and mCHH (**c**,**d**) in GLTs, wGLTs and other TEs (n=73, 232 and 3,598, respectively) in WT, *cc*, *sss* and F1 (**a**,**c**) and in WT, *h2a.w*, *wcc*, *wsss* and *w*F1 (**b**,**d**). The data of WT, *cc*, *sss* and F1 are from previously published results (GSE14875318).

### Fig.S3

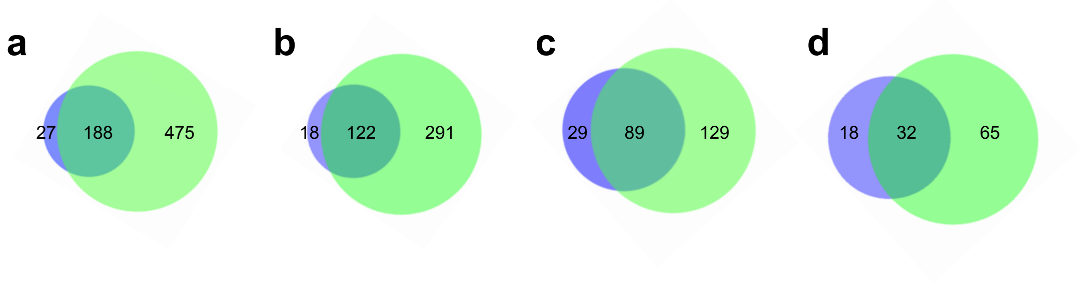


***h2a.w*** **mutation reduces the efficiency of mCH recovery in hundreds of TEs with fragments and methylated protein-coding genes**

**a** Venn diagram showing overlap between TEs without mCHG recovery in F1 (blue) and in *w*F1 (green). TEs including TE fragments with mCHG (> 0.2) in both WT and *h2a.w*, with low mCHG (< 0.1) in all of *cc*, *sss*, *wcc* and *wsss* (n=6,405), and without mCHG recovery in F1 (F1/WT< 0.1) or in *w*F1 (*w*F1/*h2a.w* <0.1) were extracted and shown. **b** Venn diagram showing overlap between TEs without mCHH recovery in F1 (blue) and in *w*F1 (green). TEs including TE fragments with mCHH (> 0.05) in both WT and *h2a.w*, with low mCHH (< 0.03) in all of *cc*, *sss*, *wcc* and *wsss* (n= 3,232), and without mCHH recovery in F1 (F1/WT< 0.1) or in *w*F1 (*w*F1/*h2a.w* <0.1) were extracted and shown. **c,d** The same as **a** and **b**, respectively, except for showing the overlap between protein-coding genes without mCH recovery.

### Fig.S4

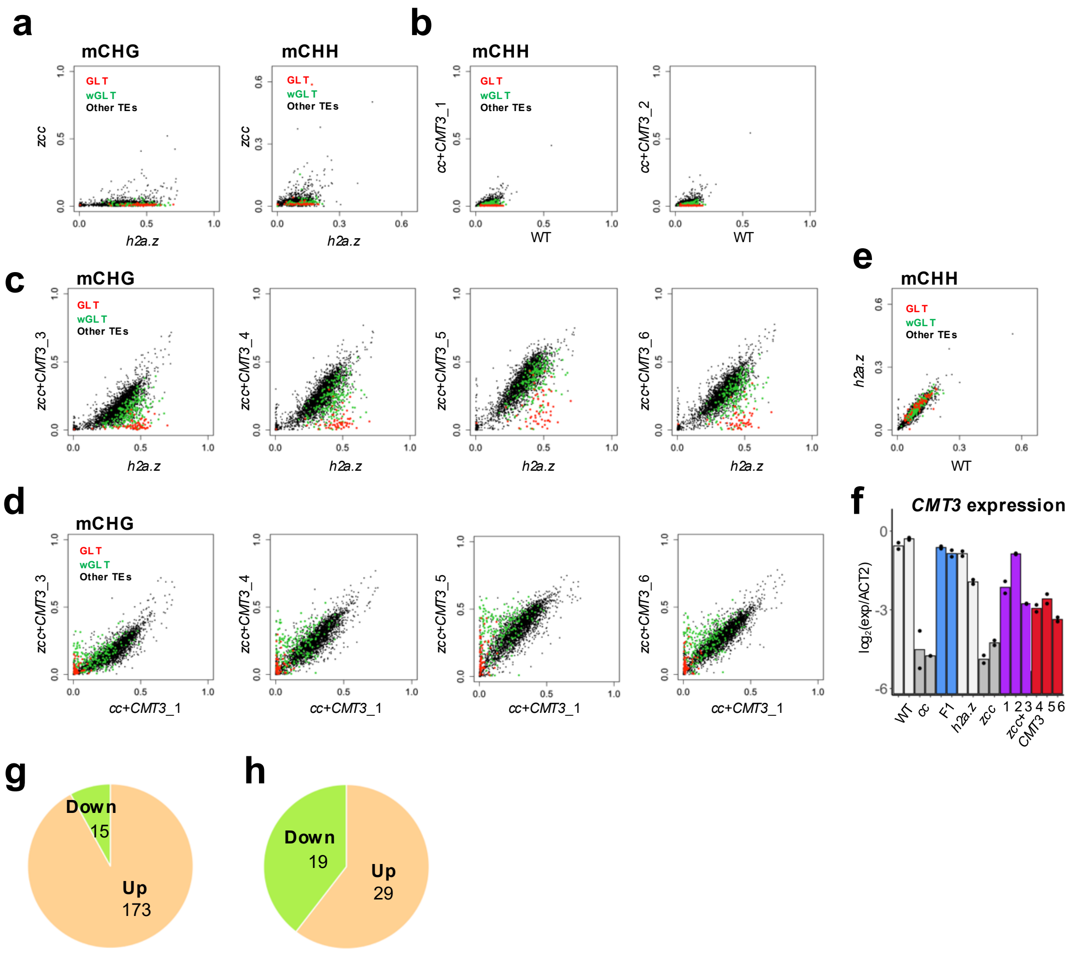


***h2a.z* mutation enables mCHG recovery in GLTs and wGLTs in *zcc+CMT3* plants**

**a** mCHG (left) and mCHH (right) level of each TE in *zcc* compared to *h2a.z*. GLTs are indicated in red, wGLTs in green, and the other TEs in black. **b** mCHH level of each TE in *cc+CMT3* compared with WT. **c**,**d** mCHG level of each TE in *zcc+CMT3* compared with *h2a.z* (**c**) and with *cc+CMT3*_1 (**d**). **e** mCHH level of each TE in *h2a.z* compared with WT. **f** The results of RT-qPCR for the transcripts of *CMT3* relative to *ACT2* in WT, *cc*, F1, *h2a.z*, *zcc* and six individuals of *zcc+CMT3*s. Two biological replicates were analyzed. Although the overall efficiency of mCH recovery varied among individual *z*T1 replicates (Fig. 2e, S4c), a correlation was not identified between the *CMT3* transcript levels and mCHG recovery. **g**,**h** Pie chart showing the number of TE fragments (**g**) and protein-coding genes with mCHG >0.05 in WT (**h**) with statistically changes in mCHG in *zcc+CMT3*_1-3.

### Fig.S5

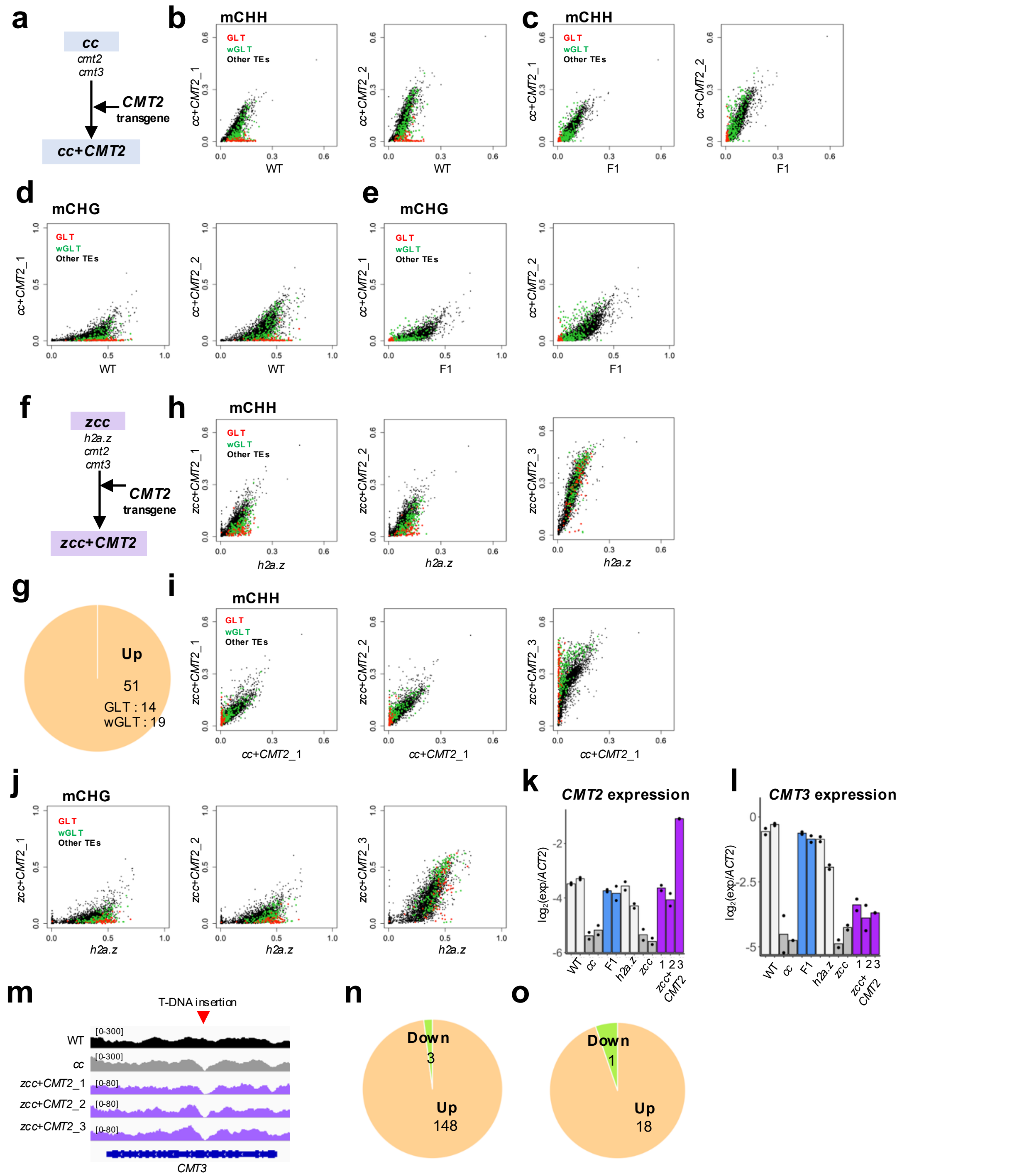


***CMT2* gene complementation in the presence of *H2A.Z* and *h2a.z* mutant background**

**a** Experimental design in the presence of *H2A.Z*. *CMT2* gene was complemented by the transformation of *CMT2* transgene into *cmt2 cmt3*: *cc* in the presence of *H2A.Z*. **b,c** Comparison of mCHH levels for each TE in the two *cc+CMT2* individuals with WT (**b**) and with F1 (**c**). GLTs are indicated in red, wGLTs in green, and the other TEs in black. **d,e** mCHG levels for each TE in the two *cc+CMT2* individuals with WT (**d**) and with F1 (**e**). **f** Experimental design in the absence of *H2A.Z*. *CMT2* gene was complemented by the transformation of *CMT2* transgene into *h2a.z cc* mutant (*hta8 hta9 hta11 cmt2 cmt3*: *zcc*). **g** Pie chart showing the number of TEs with statistically changes in mCHH in *zcc+CMT2* plants compared to *cc+CMT2* plants. The mCHH levels were compared between two individuals of *cc+CMT2* plants and three individuals of *zcc+CMT2* plants. **h,i** mCHH levels of each TE in the *zcc+CMT2* individuals compared with *h2a.z* (**h**) and with *cc+CMT2* (**i**). GLTs are indicated in red, wGLTs in green, and the other TEs in black. **j** mCHG levels of each TE in three *zcc+CMT2* individuals compared with *h2a.z*. **k,l** The results of RT-qPCR for the transcripts of *CMT2* (**k**) and *CMT3* (**l**) relative to *ACT2* in WT, *cc,* F1, *h2a.z*, *zcc* and three individuals of *zcc+CMT2*. Two biological replicates were analyzed for WT, *cc,* F1, *h2a.z* and *zcc*. *CMT2* transcripts were comparable to WT and F1 in *zcc+CMT2*_1 and 2 individuals, which showed comparable mCHH recovery to those in *cc+CMT2*s. In *zcc+CMT2*_3, however, *CMT2* transcript was approximately four times higher than WT, which was associated with a notable increase in overall mCHH recovery and evident increase in mCHG (Fig. S5h-j). Low *CMT3* transcripts was confirmed as in *zcc* and other *zcc+CMT2* plants (**l**), implying that *CMT2* overexpression can induce hypermethylation at CHH sites, as well as notable methylation at CHG sites, as reported previously[1,2]. **m** Coverage of the BS-seq data for WT, *cc* and three *zcc+CMT2* individuals at *CMT3* locus. The red arrow indicates the T-DNA insertion site. Homozygous mutation of *CMT3* was confirmed by sharp decrease in BS-seq coverage around T-DNA insertion site, besides the conventional genotyping by PCR. **n,o** Pie chart showing the number of TE fragments (**n**) and protein-coding genes with mCHH >0.03 in WT (**o**) with statistically changes in mCHH in *zcc+CMT2*.

### Fig.S6

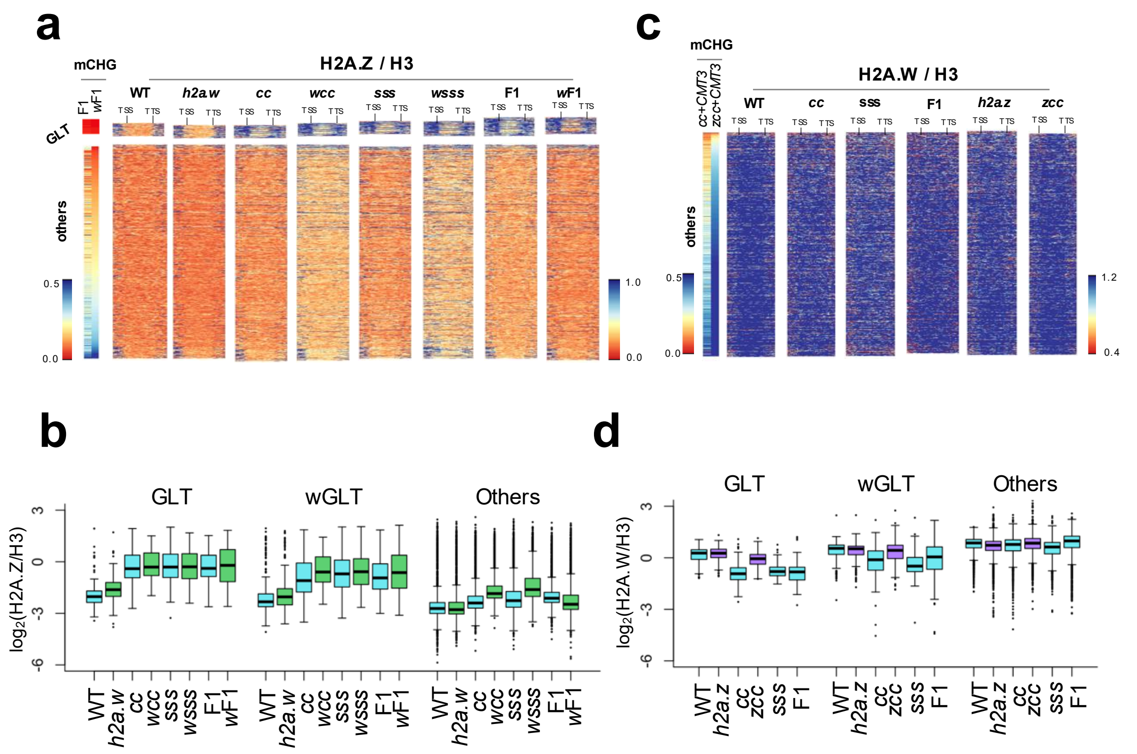


***h2a.w* mutation induces ectopic H2A.Z in wGLTs and *h2a.z* mutation suppresses H2A.W reduction in GLTs and wGLTs**

**a** Heatmaps of H2A.Z over H3 within GLTs and other TEs except for GLTs and wGLTs and their flanking regions (2 kb) in WT, *h2a.w*, *cc*, *wcc*, *sss*, *wsss*, F1 and *w*F1. TEs with length > 1000 bp and mCHG in both WT and *h2a.w* > 0.1 are analyzed (GLTs: n=64, others: n=2,330). Each TE within each group is sorted according to the mCHG levels in *w*F1. **b** Enrichments of H2A.Z over H3 in the indicated plants in GLTs, wGLTs and the other TEs. **c** Heatmaps of H2A.W over H3 within TEs except for GLTs and wGLTs and their flanking regions (2 kb) in WT, *cc, sss,* F1, *h2a.z* and *zcc*. TEs with length > 1000 bp and mCHG in both WT and *h2a.z* > 0.1 are analyzed (n=1,694). Each TE within each group is sorted according to the mCHG levels in *zcc+CMT3*_1. **d** Enrichment of H2A.W over H3 in the indicated plants in GLTs, wGLTs and the other TEs. TEs with mCHG in WT > 0.1 are analyzed.

### Fig.S7

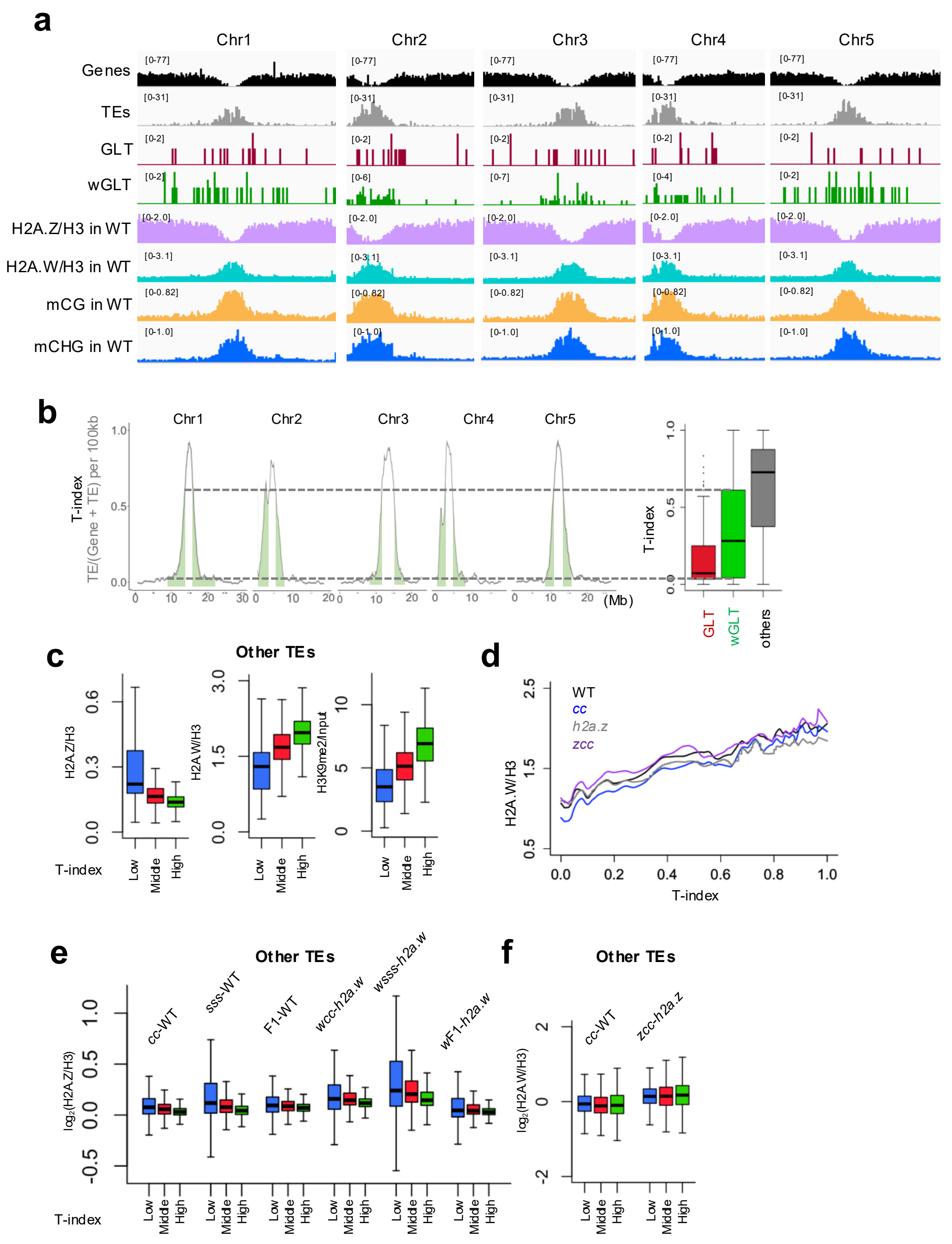


**T-index around TEs affects the localization of H2A.Z, H2A.W and H3K9me2**

**a** The distributions of GLTs and wGLTs on all 5 chromosomes of Arabidopsis genome. The numbers of TEs (gray) and normal protein-coding genes (black), and the average enrichments of H2A.Z over H3 (purple), H2A.W over H3 (green), mCG (orange), andmCHG (blue) in WT in a 100 kb bin. **b** T-index on all 5 chromosomes, comparing with that around GLTs (red), wGLTs (green) and other TEs (gray) (for 50 kb each side, 100 kb in total) indicated as Fig. 4d. **c** The enrichments of H2A.Z/H3 (left), H2A.W/H3 (middle) and H3K9me2/input (right) in WT for other TEs than with GLTs and wGLTs (n=3,598), which are divided into three groups with low T-index (<0.33), middle T-index (between 0.33 and 0.67), and high T-index (>0.67). **d** LOESS curves for the enrichment of H2A.W/H3 in TEs against T-index. TEs with mCHG in both WT and *h2a.z* (> 0.1, n=3,354) are analyzed. **e** Changes in H2A.Z/H3 enrichment compared to WT or to *h2a.w* for other TEs than GLTs and wGLTs. Other TEs are divided into three groups according to the T-index shown as in **c**. **f** Changes in H2A.W/H3 enrichment compared to WT or to *h2a.z* for other TEs than GLTs and wGLTs shown as in **c**.

### Fig.S8

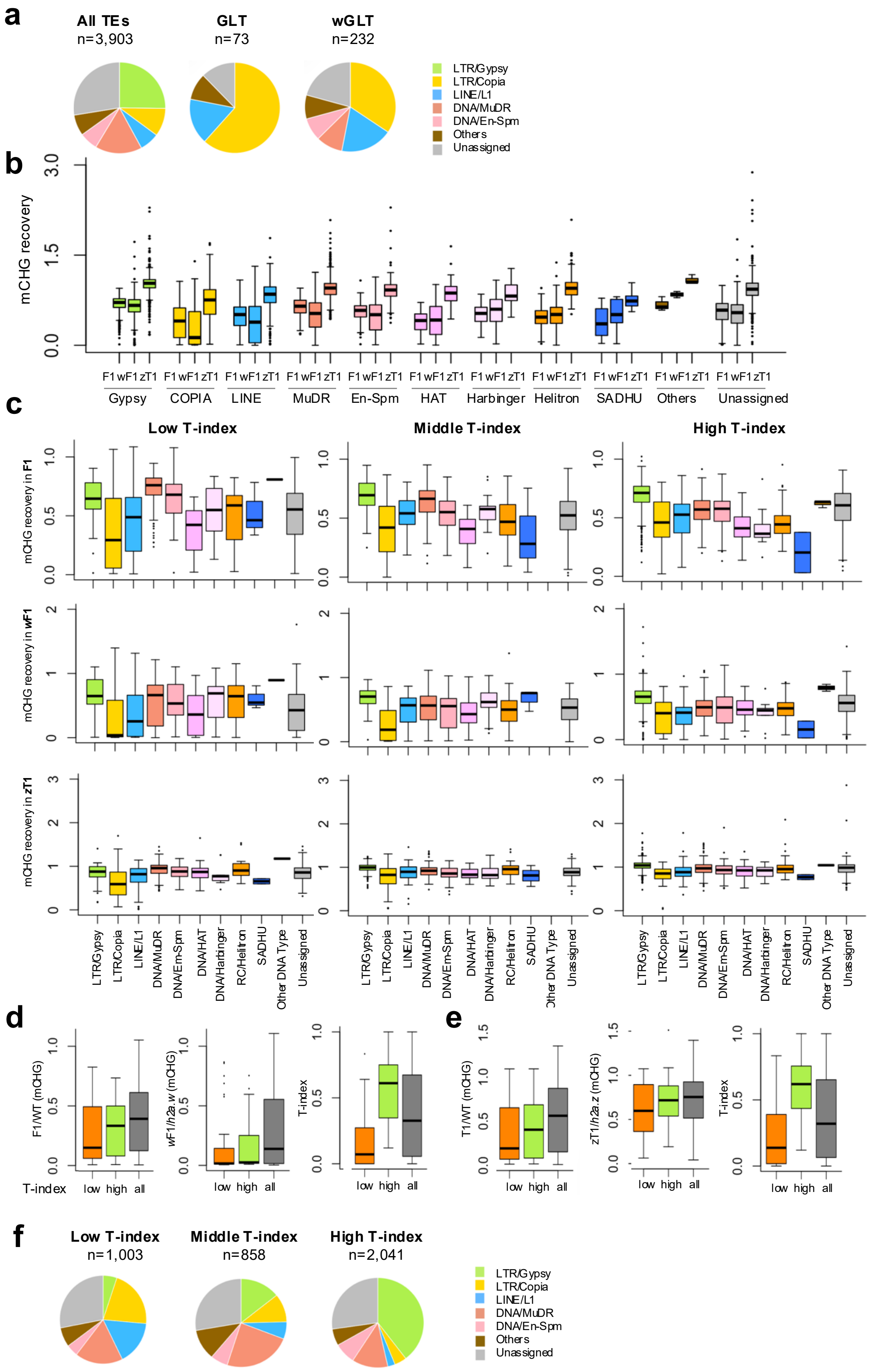


**Both T-index and TE family affect the efficiency of mCH recovery**
**a** Pie chart of the proportion of TE family in all TEs, GLTs and wGLTs. **b** TE families are compared for mCHG recovery in F1 (F1/WT), *w*F1 (*w*F1/*h2a.w*) and *z*T1 (*zcc+CMT3*/*h2a.z*). **c** TE families are divided into three according to the T-index (low, T-index<0.33; middle, 0.33<=T-index<0.67; high, T-index>=0.67) and are compared for mCHG recovery in F1 (F1/WT), *w*F1 (*w*F1/*h2a.w*) and *z*T1 (((*zcc+CMT3*_1+ *zcc+CMT3*_2 + *zcc+CMT3*_3)/3) / *h2a.z*). **d,e** The phylogenetically close pairs of *COPIAs* (see methods for detail) were divided into two groups with relatively low (orange) and high (green) T-index, then compared for the mCHG recovery in F1 and *w*F1 (n=29 each, **d**), or in T1 and *z*T1 (n=23 each, **e**), as well as for the T-index. For comparison, all *COPIAs* (grey, n=257) are shown for each plot. **f** Pie chart of the proportion of TE family in TEs with low, middle and high T-index (low, T-index<0.33; middle, 0.33<=T-index<0.67; high, T-index>=0.67).

### Fig.S9

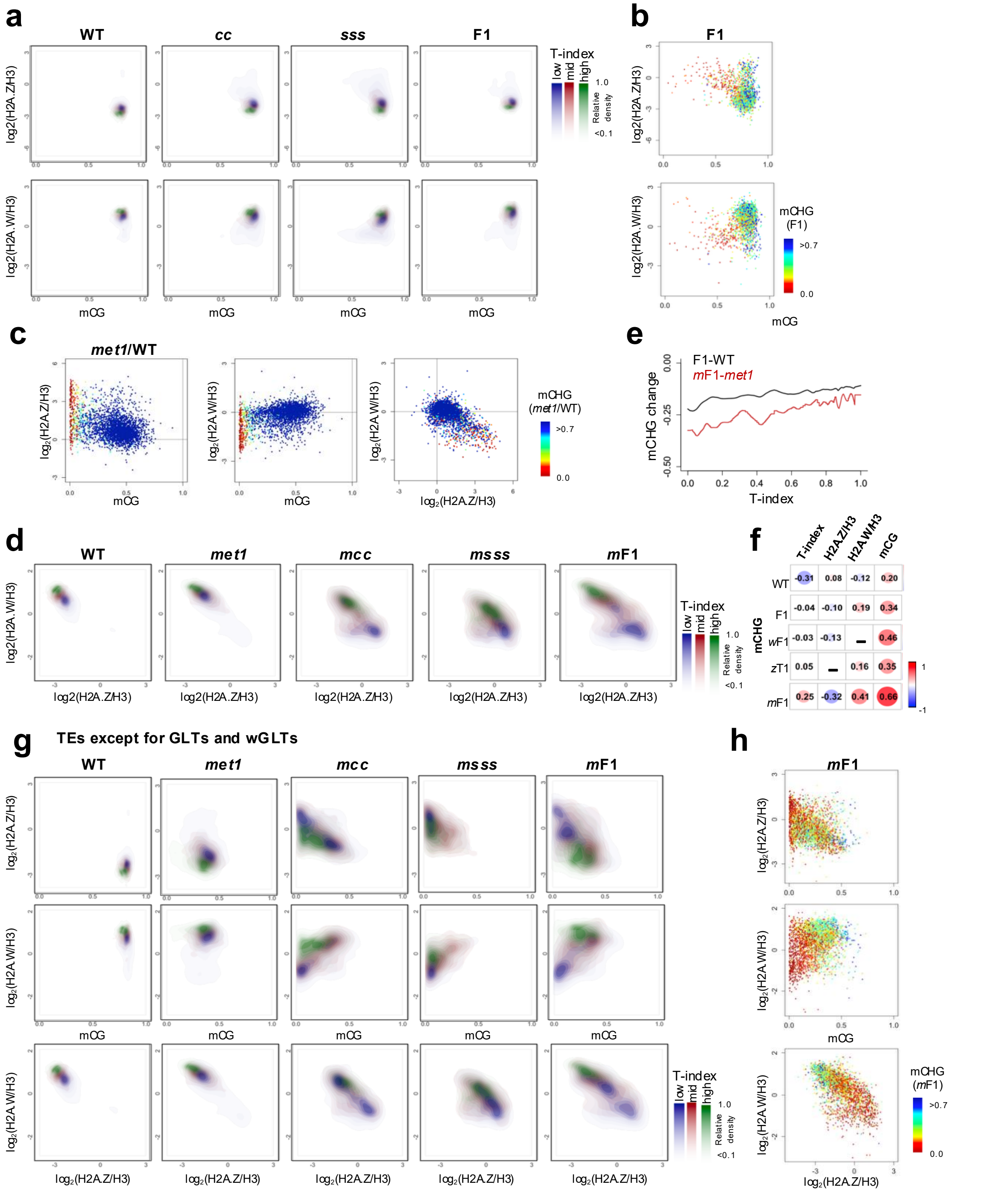


**mCH recovery and localization of H2A.W and H2A.Z are dependent on chromosomal environments**
**a** Density plots for the enrichments of H2A.Z/H3 (upper) and H2A.W/H3 (lower) against mCG levels, comparing among the three groups of TEs (blue for low T-index (< 0.33); red for middle T-index (0.33 ≤ T-index < 0.67), green for high T-index ≥ 0.67). **b** Scatter plots comparing between mCG and H2A.Z/H3 (upper) or H2A.W/H3 (lower) in TEs in F1. TEs with mCHG in WT > 0.1 (n=3,538) are analyzed. Each TE is colored according to mCHG level in F1. **c** Each TE is compared among mCG, H2A.Z/H3, and H2A.W/H3 in *met1* relative to WT. To avoid division by values near zero, TEs with less than 0.1 in WT for any of mCHG, mCG, H2A.W/H3, and H2A.Z/H3 are excluded from the analysis (n=3,237). **d** Density plots for the enrichments of H2A.Z/H3 against H2A.W/H3, comparing among the three groups of TEs as indicated in **a**. **e** LOESS curves for the mCHG change in TEs between F1 and WT (black), and between *m*F1 and *met1-1* (red) against T-index. TEs with mCHG in both WT and *met1* (> 0.1, n=3,212) are analyzed. Data from GSE148753 and GSE181896 were re-analyzed. **f** Spearman's rank correlation coefficient between mCHG levels and T-index, H2A.Z/H3, H2A.W/H3 and mCG levels in TEs. Both the color and size of circles represent spearman's rank correlation coefficient. **g** Density plots for the enrichments of H2A.Z/H3 (upper) and H2A.W/H3 (middle) against mCG levels and H2A.Z/H3 against H2A.W/H3 (bottom), comparing among the three groups of TEs except for GLTs and wGLTs as indicated in **a**. **h** Scatter plots comparing between mCG and H2A.Z/H3 (upper) or H2A.W/H3 (lower) in TEs in *m*F1. TEs except for GLTs and wGLTs with mCHG in both WT and *met1* > 0.1 (n=2,934) are analyzed. Each TE is colored according to mCHG level in *m*F1.

### Fig.S10

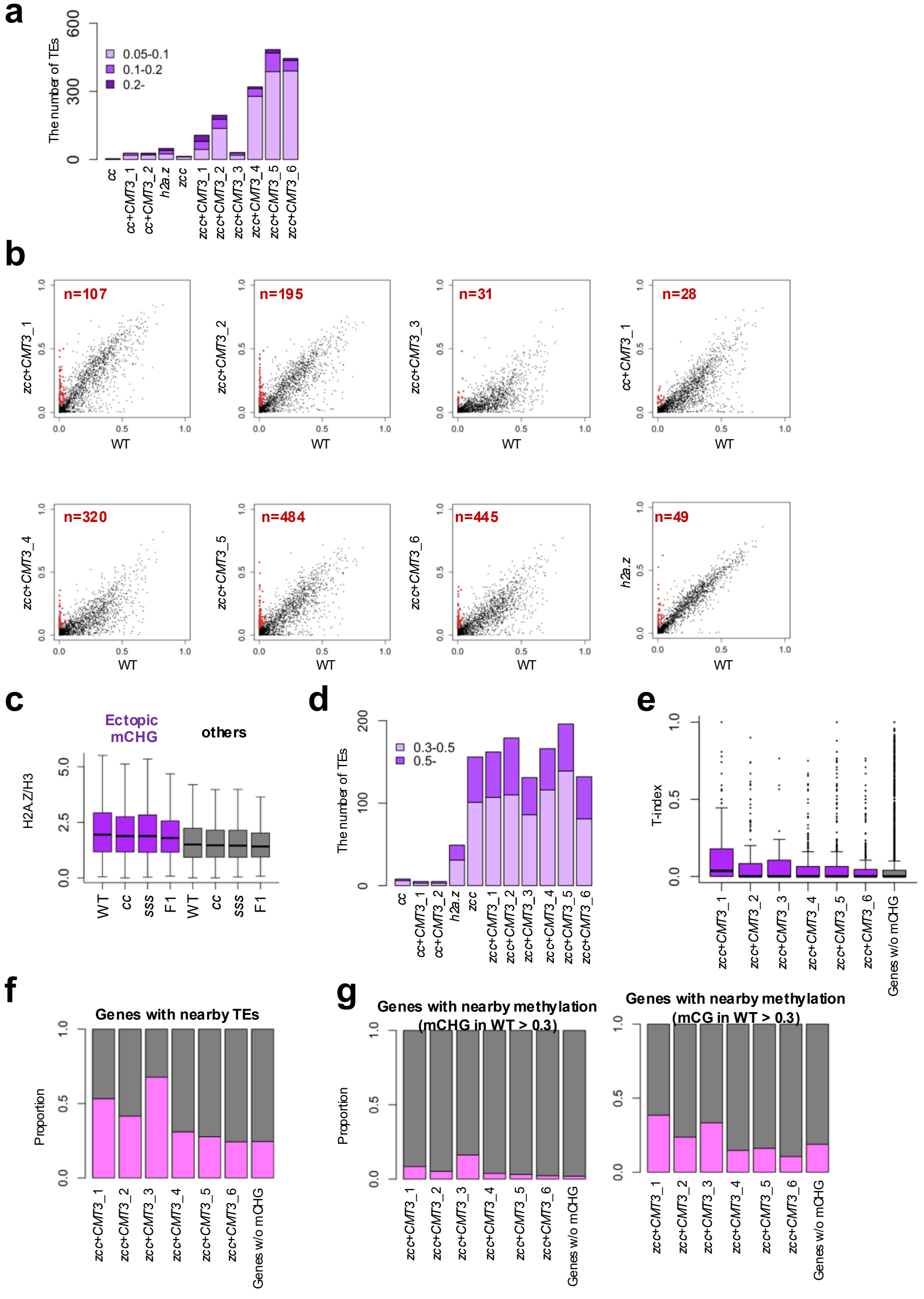


**mCH/H3K9me reconstitution in *h2a.z* induces ectopic heterochromatin in protein-coding genes**

**a** The numbers of protein-coding genes with ectopic mCHG. Protein-coding genes without mCHG in WT (WT mCHG< 0.05) and with higher mCHG in the indicated genotypes than in WT with difference 0.05-0.1, 0.1-0.2 or above 0.2 were counted. **b** Scatter plots for mCHG levels of each protein-coding gene in each *z*T1 individual, *cc+CMT3*_1, and *h2a.z* compared with WT. Protein-coding genes without mCHG in WT (< 0.05) and each *zcc+CMT3*-WT, *cc+CMT3*_1-WT or *h2a.z*-WT (mCHG) >0.05 are indicated in red. **c** H2A.Z enrichments over H3 in protein-coding genes (mCHG in WT < 0.05) in WT, *cc*, *sss* and F1. The protein-coding genes were divided into those with ectopic mCHG > 0.05 in at least one *z*T1 individual (purple) and the others (gray). Outliers are excluded. **d** The numbers of protein-coding genes with ectopic mCG. Protein-coding genes without mCHG in WT (WT mCHG< 0.05) and with higher mCG in the indicated genotypes than in WT with difference 0.3-0.5, and above 0.5 were counted. **e** T-index around protein-coding genes with ectopic mCHG in WT <0.05 and mCHG > 0.1 in each *z*T1 individual (purple) and the other genes with mCHG <0.05 in WT (gray). **f** The proportion of protein-coding genes with TEs or TE fragments in their flanking regions (±500 bp) in the mCHG-gained genes in each *z*T1 and in the other genes without mCHG (WT mCHG<0.05) shown in pink. **g** The proportion of protein-coding genes with DNA methylated franking regions (±500 bp). The franking regions of mCHG-gained genes in each *z*T1 and the other genes without mCHG (WT mCHG<0.05) were analyzed for DNA methylation, and the proportion of the genes within the genes in WT <0.05 with mCHG >0.3 (left) or with mCG >0.3 (right) are shown in pink.

### Table S1

**Supplementary Table 1. Spearman correlation with H3K9me2 and mCH recovery**

Spearman correlation between each of the distance from centromeres to TEs, TE density per 100kb around TEs, Gene density per 100kb around TEs, and T-index around TEs with H3K9me2/Input in WT and mCHG recovery in F1 (F1/WT). TEs with mCHG in WT >0.1 were used for the analysis.

| Factors | H3K9me2/Input | mCH recovery (F1/WT) |
|---|---|---|
| The distance from centromeres | -0.487 | -0.0851 |
| TE density per 100kb | 0.514 | 0.0659 |
| Gene density per 100kb | -0.595 | -0.132 |
| T-index per 100kb | 0.608 | 0.136 |

### Supplementary References

1. Stroud, H. et al. Non-CG methylation patterns shape the epigenetic landscape in Arabidopsis. *Nat. Struct. Mol. Biol.* **21**, 64–72 (2014).

2. Jiang, J. et al. Substrate specificity and protein stability drive the divergence of plant-specific DNA methyltransferases. *Sci. Adv.* **10**, eadr2222 (2024).
